# Cell dynamics underlying oriented growth of the *Drosophila* wing imaginal disc

**DOI:** 10.1101/140038

**Authors:** Natalie A. Dye, Marko Popović, Stephanie Spannl, Raphaël Etournay, Dagmar Kainmüller, Eugene W. Myers, Frank Jülicher, Suzanne Eaton

## Abstract

Quantitative analysis of the dynamic cellular mechanisms shaping the *Drosophila* wing during its larval growth phase has been limited, impeding our ability to understand how morphogen patterns regulate tissue shape. Such analysis requires imaging explants under conditions that maintain both growth and patterning, as well as methods to quantify how much cellular behaviors change tissue shape. Here, we demonstrate a key requirement for the steroid hormone 20-hydroxyecdysone (20E) in the maintenance of numerous patterning systems *in vivo* and in explant culture. We find that low concentrations of 20E support prolonged proliferation in explanted wing discs in the absence of insulin, incidentally providing novel insight into the hormonal regulation of imaginal growth. We use 20E-containing media to directly observe growth and apply recently developed methods for quantitatively decomposing tissue shape changes into cellular contributions. We discover that while cell divisions drive tissue expansion along one axis, their contribution to expansion along the orthogonal axis is cancelled by cell rearrangements and cell shape changes. This finding raises the possibility that anisotropic mechanical constraints contribute to growth orientation in the wing disc.

## INTRODUCTION

The *Drosophila* larval wing imaginal disc is a powerful model system in which to study the integration of diverse types of regulatory cues for tissue growth. The wing disc is a relatively flat epithelial sac that grows to about 50,000 cells during larval stages of development. As it grows, the rapidly proliferating cells on one side (the wing pouch) become pseudostratified, while cells on the other side become squamous. At pupariation, the wing pouch everts and flattens to assume the adult wing shape (Waddington 1940).

The size and shape of the wing are largely, but not entirely, determined during the larval growth phase. At this stage, signaling centers located at the anterior-posterior (AP) and dorsal-ventral (DV) compartment boundaries not only establish patterns of gene expression but also promote growth (Beira & Paro 2016; Hartl & Scott 2014). Hedgehog produced in posterior cells signals across the AP boundary to stabilize the transcriptional activator Ci155 and induce the expression of different target genes (i.e., Engrailed and Decapentaplegic (Dpp)) at different distances (reviewed in (Hartl & Scott 2014)). Dpp is itself a secreted signaling molecule that establishes a bidirectional gradient anteriorly and posteriorly, promoting patterned gene expression through graded phosphorylation of SMAD transcription factors (reviewed in (Affolter & Basler 2007)). At the DV boundary, Wingless (Wg) expression and Notch signaling are maintained by a positive feedback loop and together pattern the DV axis (Micchelli & Blair 1999; Micchelli et al. 1997; Rulifson et al. 1996). Signaling from the AP and DV organizers is required for wing disc growth, even though cell proliferation is not especially concentrated near these regions (Schwank et al. 2011; González-Gaitán et al. 1994; Milán et al. 1996).

The orientation of tissue growth is strikingly non-uniform: marked clones strongly elongate along the proximo-distal (PD) axis of the adult wing (González-Gaitán et al. 1994; Resino et al. 2002; Baena-López et al. 2005; Mao et al. 2013; Worley et al. 2013; Heemskerk et al. 2014). In central regions of the larval wing pouch, the adult PD axis is generally orthogonal to the DV boundary. Signaling at organizer regions influences the growth orientation by establishing tissue-wide patterns of two planar cell polarity (PCP) systems, Fat and Core, with distinct cortical complexes that are polarized in the epithelial plane (Aw & Devenport 2016). Both systems develop the same global pattern of planar polarity, which is oriented with respect to the AP and DV boundaries and presages the orientation of growth (Rogulja et al. 2008; Ambegaonkar et al. 2012; Sagner et al. 2012; Brittle et al. 2012). Perturbing Fat PCP or dominantly reversing the orientation of Core PCP both disrupt wing size and shorten it in the PD axis (Bryant et al. 1988; Clark et al. 1995; Mao et al. 2006; Merkel et al. 2014).

A complete description of how the PCP patterns direct oriented tissue growth is still missing. PD-oriented cell divisions clearly contribute to oriented tissue growth and are disturbed by genetic disruption of the Fat pathway (Baena-López et al. 2005; Mao et al. 2011). However, quantitatively, it is still unknown the extent to which oriented cell divisions or other dynamic behaviors (T1 rearrangements, cell shape changes or extrusions) determine the tissue growth pattern. Analytical tools now exist to fully describe tissue growth from cellular dynamics (Merkel et al. 2017; Etournay et al. 2015; Etournay et al. 2016; Guirao et al. 2015), but such analysis demands long-term time-lapse imaging at high spatial and temporal resolution. *In vivo* imaging has an insufficient time resolution (Heemskerk et al. 2014), necessitating the use of *ex vivo* culture. The achievement of prolonged *ex vivo* wing disc growth has proven difficult, even though the dramatic eversion process that imaginal tissues undergo at the onset of pupariation can be successfully replicated in culture (Fristrom et al. 1973; Milner 1977; Aldaz et al. 2010). Importantly, in order to properly study the cellular dynamics underlying oriented growth, we require a culture condition that maintains the signaling from AP/DV organizers and global PCP patterns, and the consequences for culture on these systems is completely unknown.

Insulin signaling regulates animal size in response to nutrition, and bovine insulin has been widely used in culture to stimulate growth of explanted discs (Zartman et al. 2013; Mao et al. 2011; Mao et al. 2013; Legoff et al. 2013; Heller et al. 2016; Tsao et al. 2016). However, proliferation in insulin-cultured discs arrests within a few hours (Handke et al. 2014; Tsao et al. 2016); thus clearly other signals are required for long term growth. One such missing signal could be ecdysteroids, a family of steroid hormones produced by the ring gland in arthropods with well characterized functions in regulating major developmental transitions (Kozlova & Thummel 2000). A peak of 20-hydroxyecdysone (20E) and related hormones at the larval to pupal transition induces pupariation and imaginal disc eversion (Fristrom et al. 1973; Milner 1977; Aldaz et al. 2010). This peak is preceded by a smaller elevation in hormone levels (approximately 25-fold lower concentration) that starts in the mid-third larval instar (Kozlova & Thummel 2000; Lavrynenko et al. 2015). Several lines of evidence now suggest that these lower concentrations promote imaginal growth and development during larval stages (Bodenstein 1943; Brennan et al. 1998; Brennan et al. 2001; Mirth et al. 2009; Delanoue et al. 2010; Mitchell et al. 2013; Herboso et al. 2015).

Here, we show that physiologically low levels of 20E, in the absence of insulin, significantly extend the proliferation of explanted wing discs over insulin alone. Furthermore, we show with transcriptome sequencing that 20E is required in culture for the normal expression of numerous genes involved in wing patterning, whereas insulin promotes a strong but transient anabolic growth response. Genetic perturbations *in vivo* confirm that 20E has widespread effects on morphogen signaling and PCP in the larval wing, highlighting the importance of including 20E in culture media for replicating *in vivo* growth patterns. We exploit this improved system to perform live imaging and apply recent tools for decomposing tissue growth into contributions from each type of dynamic cell behavior (Merkel et al. 2017; Etournay et al. 2015; Etournay et al. 2016). Our analysis indicates that divisions alone are insufficient to quantitatively account for the anisotropy of tissue growth, as previously thought; cell rearrangements and cell shape changes play an equally important role. This work inspires new directions for exploration into the regulation of tissue size and shape in this system.

## RESULTS

### Low levels of 20E stimulate cell divisions in cultured wing discs

We directly tested the ability of low levels of 20E, with or without insulin, to maintain proliferation in explants from mid third instar larvae (96hr after egg laying (AEL)) grown at 25 °C. These discs should continue to proliferate for another 24hr *in vivo*, since under the conditions we use, pupariation occurs around 120hr AEL.

We monitored the numbers of phospho-histone H3-positive proliferating cells in freshly explanted discs, and compared them to discs cultured in the absence of hormones, or in the presence of 20nM 20E, 1µM insulin, or both hormones (Fig. 1A,B). In the absence of any hormones, the number of mitotic cells per area (proliferation index) is dramatically reduced by 4hr and negligible by 9hr (Fig. 1Ai, Bi), consistent with previous work (Zartman et al. 2013; Handke et al. 2014). In insulin, the proliferation index drops to half that of freshly explanted discs after 4hr and continues to decline thereafter (Fig. 1Aii, Bii). These indices quantitatively agree with those of previous analyses of insulin-culture discs (Handke et al. 2014).

**Figure 1:**
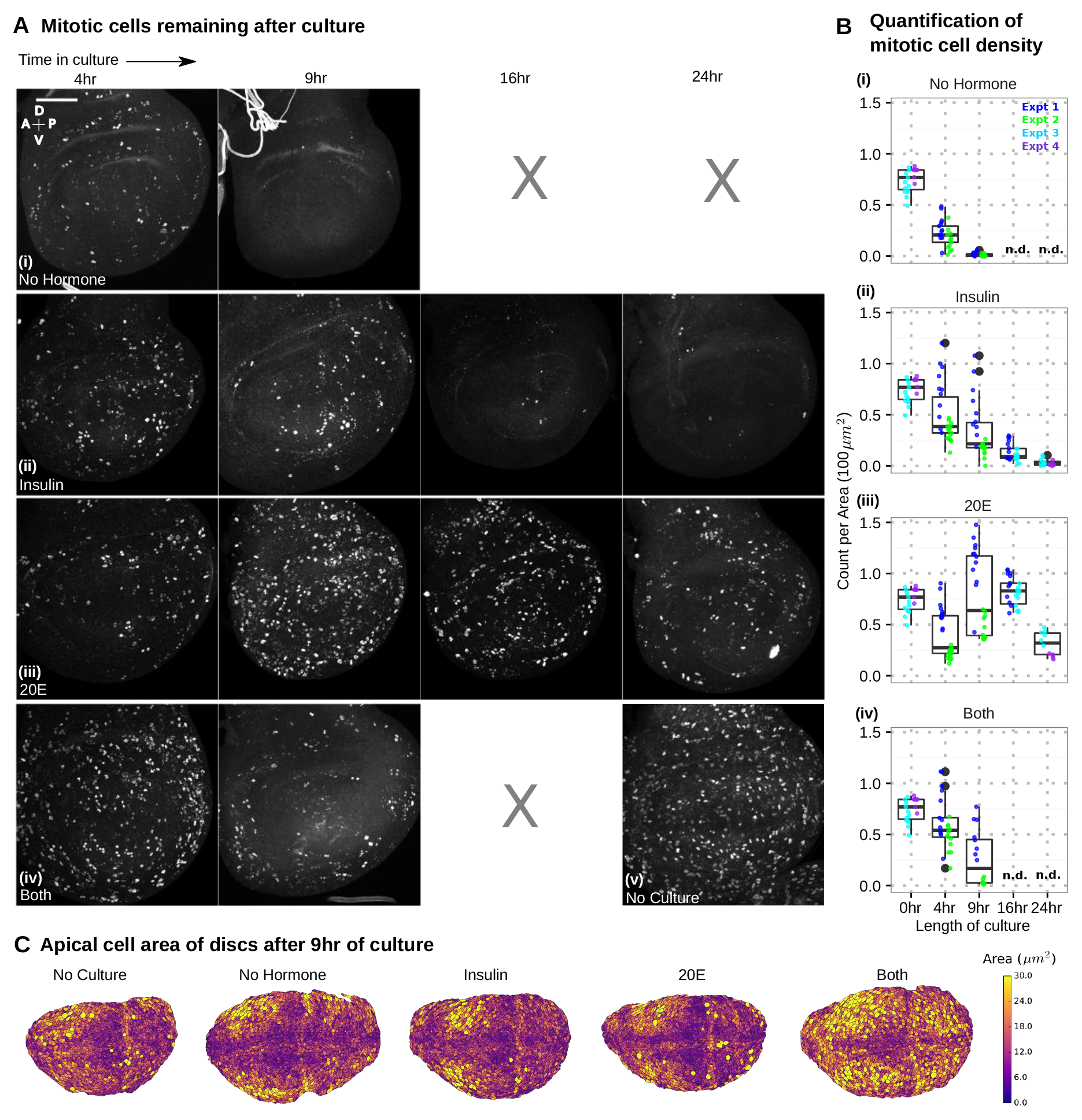
20E is sufficient to stimulate prolonged cell division in cultured wing discs. Wing discs from mid-third instar larvae (96hr AEL) were cultured for indicated times and then stained with phospho-Histone H3 to detect mitotic nuclei at the indicated timepoints. (A) Representative images and (B) their quantification. Scale bar indicates 50μm. Discs were given (i) no hormone supplement, (ii) 5μg/ml insulin, (iii) 20nM 20E or (iv) both. In (v) is an image of an uncultured disc from the beginning of the experiment (96hr AEL). In (B), we show pooled results from multiple experiments from different days. Each dot is the mean density of mitotic nuclei (# nuclei per 10x10μm^2^ area, averaged across the pouch from a single disc). Discs from experiments performed on the same day have the same color. The box plots summarize all data for each condition. Hinges correspond to the first and third quartiles; lines extend to the highest and lowest values within 1.5*IQR (inter-quartile range); outlier points are black. n.d. = not determined. (C) Apical cell area the wing pouch after 9hr of culture. Overall, the morphology is maintained during culture, except in presence of both hormones. Lowering the concentration of 20E in combination with insulin does not further prolong proliferation (See Fig. 1--Figure Supplement 1).

Strikingly, in 20E alone, the proliferation index is similar to that of insulin-cultured discs at 4hr, but instead of declining thereafter, the index actually increases (Fig. 1Aiii, Biii). From 9-16hr after explant, the same density of mitotic nuclei is found in 20E-cultured discs as in freshly explanted discs. Proliferation even extends through 24hr, albeit at a lower density than between 9-16hr. By 24hr, the overall morphology also becomes somewhat abnormal (Fig. 1—Figure Supplement 1), perhaps reflecting the fact that components of the extracellular matrix come from the fat body *in vivo* (Pastor-Pareja & Xu 2011).

The simultaneous addition of both hormones promotes proliferation mildly better than either alone for 4hr; but by 9hr, proliferation declines, and the cell area distribution becomes abnormal (Fig. 1B-C). Growth is not significantly improved by adjusting 20E concentration (Fig. 1-Figure Supplement 1). Thus, under these explant conditions, insulin antagonizes the effect of 20E on proliferation.

### Transcriptional responses to 20E and insulin in explanted wing discs

To assess the consequences for gene expression of culturing under different hormonal conditions, we sequenced the transcriptomes from freshly explanted discs, discs cultured without hormones for 4hr, and discs cultured for 4 or 9hr in either 20E or insulin alone. Relative to freshly explanted discs, the expression of 1604 genes goes down and 1409 genes goes up after 4hr of culture without hormones. Changed expression of 1911 genes (63%) could be prevented, at least transiently, by the addition of either 20E or insulin. While some genes responded to both 20E and insulin, others were hormone-specific.

Figure 2 shows the size and overlap of the hormone-responsive gene sets. Comparing the transcriptional response to insulin at 4 and 9hr reveals that it is largely transient (Fig. 2A), consistent with the decreased proliferative response to insulin by 9hr (Fig. 1Aii,Bii). In contrast, although 20E regulates a smaller subset of genes, its effects are longer lasting (Fig. 2A). Interestingly, a subset of genes can be regulated by either 20E or insulin (Fig. 2B). Maximum overlap (353 genes) occurs at time points when each hormone supports maximum proliferation (insulin at 4hr, 20E at 9hr).

**Figure 2:**
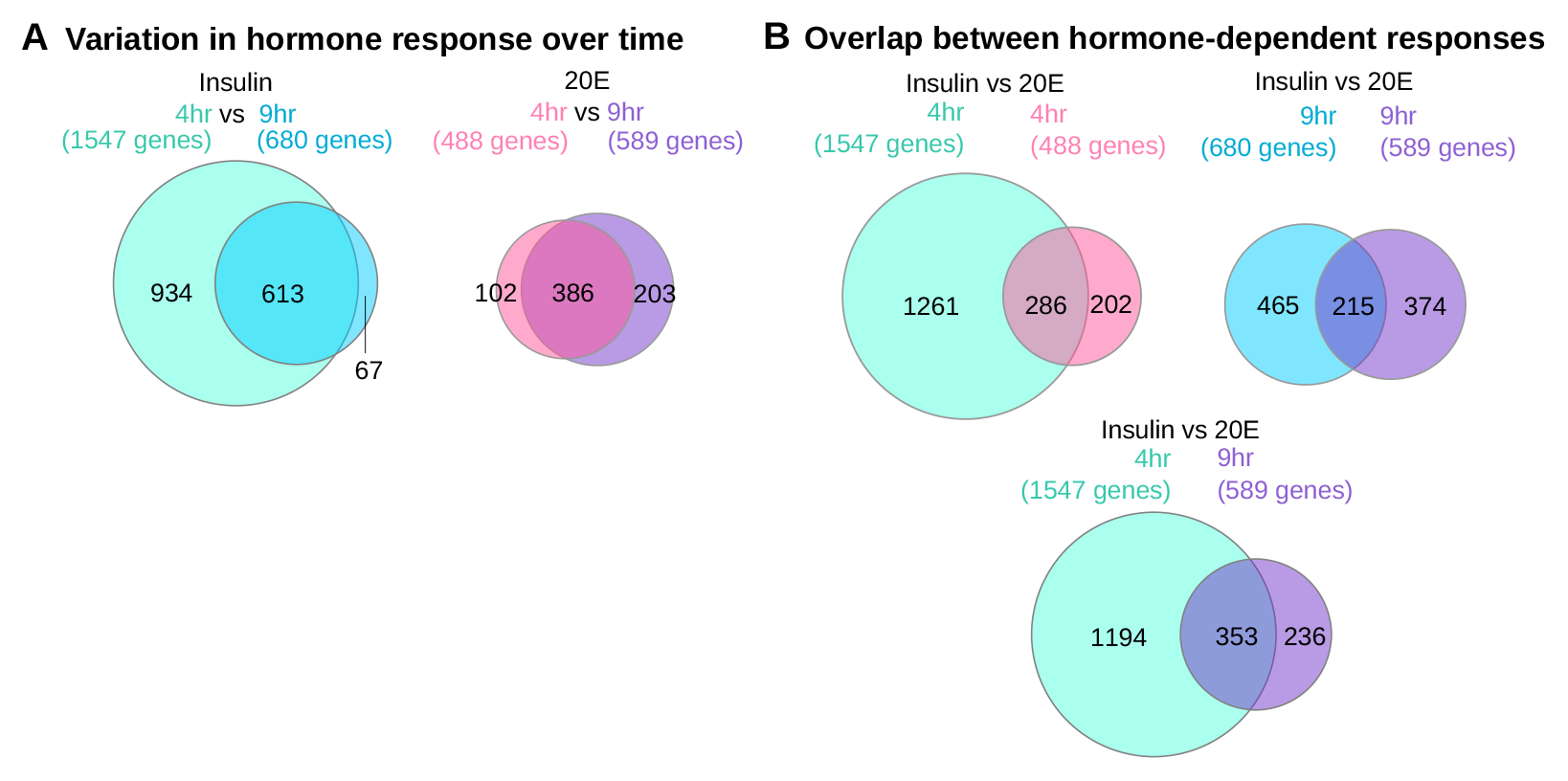
Size and overlap of the transcriptional response to 20E or insulin. Transcriptomes of wing discs cultured in either 20E or insulin were sequenced and compared to those of uncultured discs and discs cultured without any hormone. A gene was classified as hormone-dependent if its levels changed upon culture without hormone and responded in the correct direction (more similar to uncultured levels) when cultured with hormone. (A) Total number of genes that responded to 20E or insulin at two timepoints. (B) Overlap between the hormone-dependent responses at 4hr (left), 9hr (right) or when each hormone supports the most amount of proliferation (4hr for insulin, 9hr for 20E). Circles are all drawn to scale. Numbers denote the amount of genes in the overlapping or non-overlapping sets.

Insulin-induced genes are most significantly enriched in Gene Ontology (GO) terms describing DNA replication, ribosome biogenesis, translation, purine biosynthesis, and mitochondrial biogenesis (Fig. 3 and Fig. 3-Figure Supplement 1). Conversely, insulin reduces the expression of genes involved in autophagy and cell death (Fig. 3 and Fig. 3-Figure Supplement 1). Interestingly, the effect of insulin on genes in all of these GO categories is much stronger at 4hr than at 9hr of culture. This transient response to insulin must result from negative regulation of the pathway downstream of PI3K: a reporter of PI3K activation remains active even after 9hr of culture (Fig. 3-Figure Supplement 2). Thus, insulin induces a pattern of gene expression characteristic of the fed state, consistent with many previous studies (Gershman et al. 2007; Teleman et al. 2008; Li et al. 2010); however, this response is transient.

**Figure 3:**
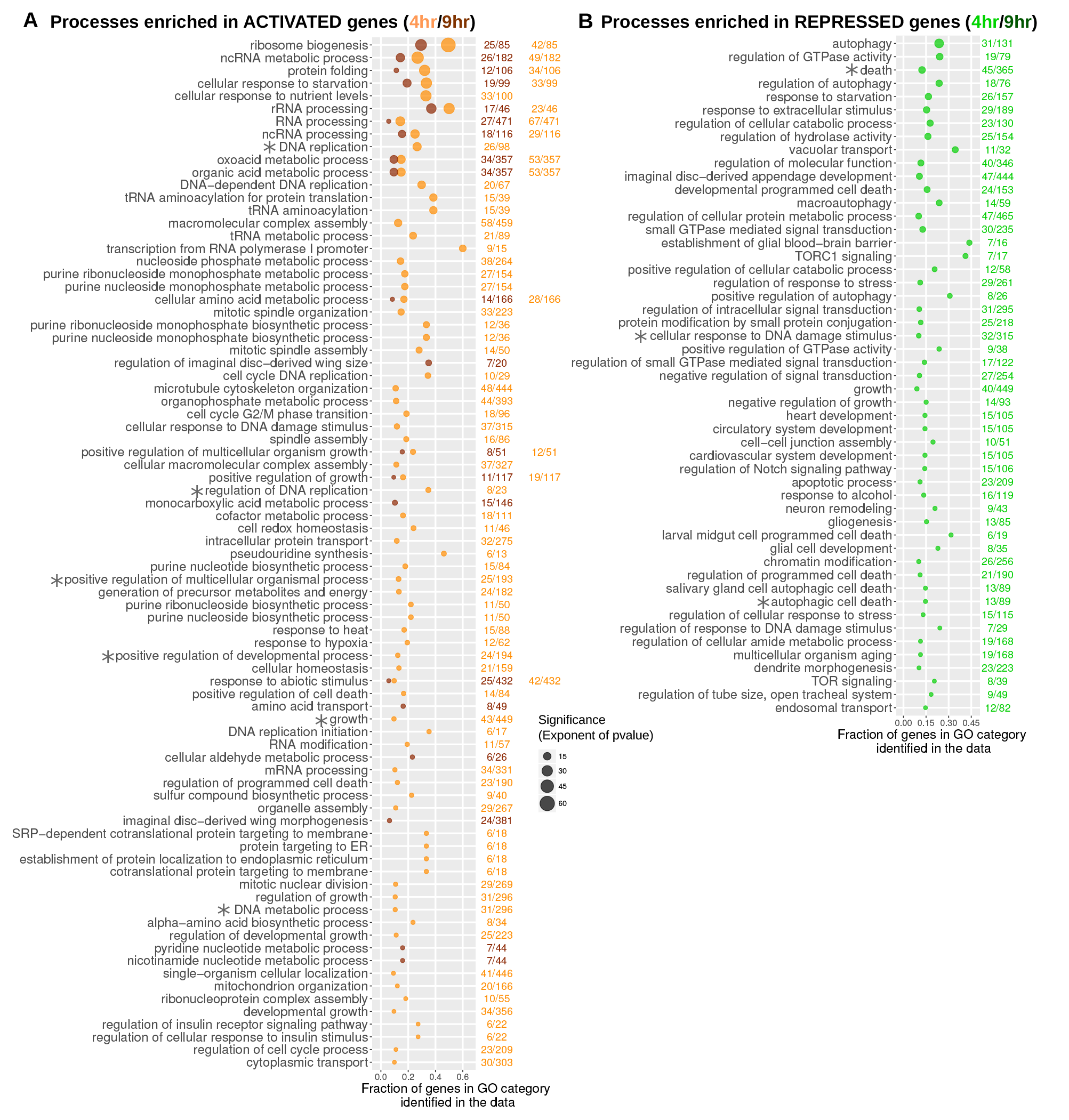
Processes that are transcriptionally regulated by insulin in cultured discs. Shown are the Biological Process (BP) Gene Ontology (GO) terms that are significantly enriched in the set of genes identified to be activated (A) or repressed (B) by insulin in cultured wing discs (simplified with a semantic similarity algorithm; threshold=0.95). In both (A) and (B), we plot the GO terms according to the fraction of its total genes that were identified in the data (# genes in the data with this GO term/total size of the GO term). The size of the point indicates the significance of enrichment (BH-corrected *p*-value; cutoff of *p* < 0.005). The color indicates either 4hr (light orange in A, green in B) or 9hr of culture (dark orange in A, dark green in B). Note that by 9hr of culture in insulin, no GO terms are enriched in the group of genes repressed by insulin (B). Asterisks indicate terms shared by both insulin and 20E. Note that many categories are only enriched (or more significantly enriched) at 4hr. This transience in the response to insulin must result from downregulation of the pathway at a point downstream of PI3K activation, since a fluorescent reporter of this enzyme’s activity indicates that it is still active for the duration of culture (See Fig 3--Figure Supplement 2).

Between the insulin and 20E-responsive gene sets, there is limited overlap of enriched GO terms (Fig. 4 and Fig. 4-Figure Supplement 1). Both hormones repress genes involved in autophagy, DNA damage response and cell death while inducing genes involved in DNA replication. Amongst the shared targets, we found a subset involved in protein synthesis and nutrient transport (Fig. 4-Figure Supplement 1C). These include 4E-BP, which was previously shown to respond to 20E *in vivo* (Herboso et al. 2015).

**Figure 4:**
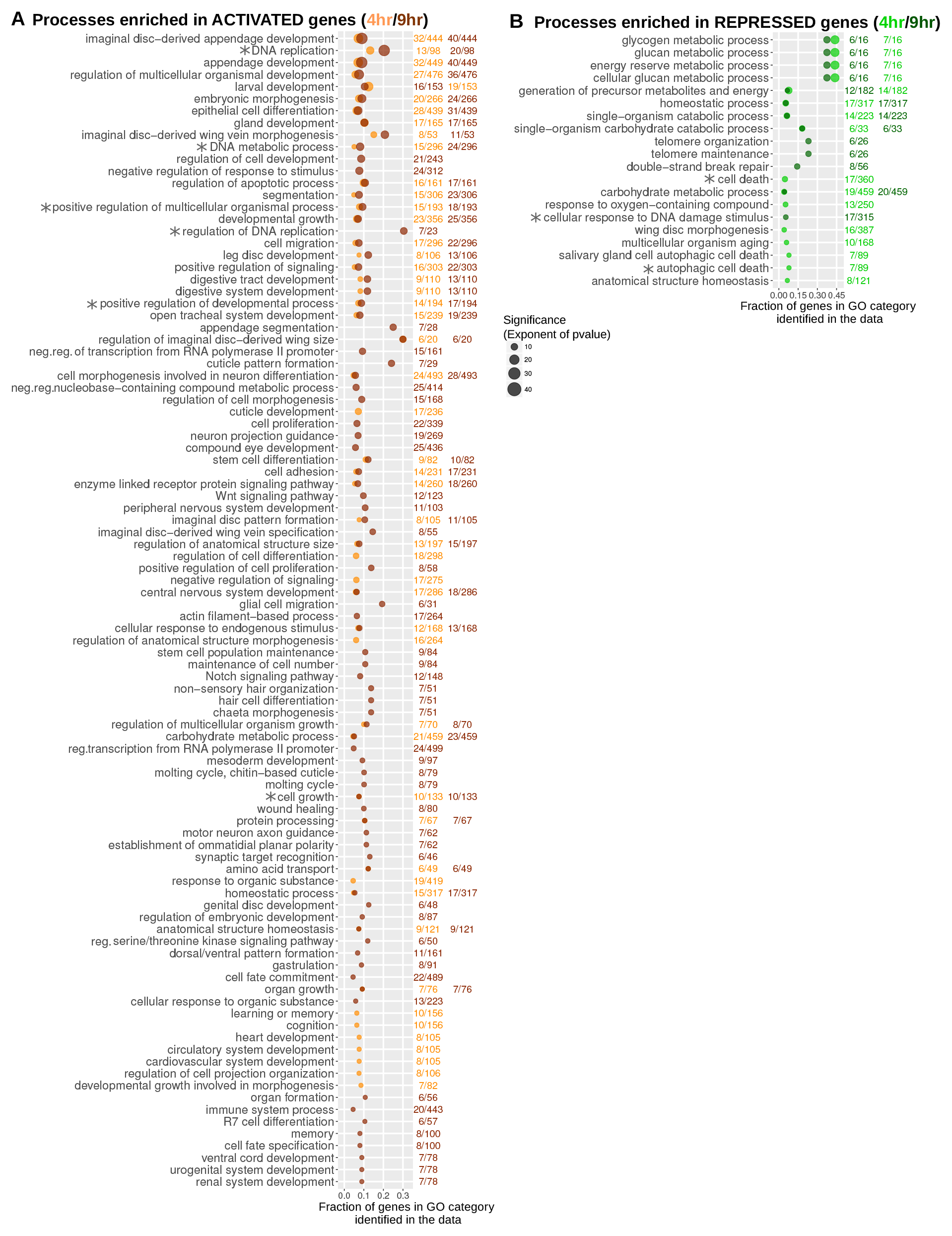
Processes that are transcriptionally regulated by 20E in cultured discs. Shown are the Biological Process (BP) Gene Ontology (GO) terms that are significantly enriched in the set of genes identified to be activated (A) or repressed (B) by 20E in cultured wing discs (simplified with a semantic similarity algorithm; threshold=0.9). In both (A) and (B), we plot the GO terms according to the fraction of its total genes that were identified in the data (# genes in the data with this GO term/total size of the GO term). The size of the point indicates the significance of enrichment (BH-corrected *p*-value; cutoff of *p* <0.005). The color indicates either 4hr (light orange in A, green in B) or 9hr of culture (dark orange in A, dark green in B). Asterisks indicate terms shared by both insulin and 20E.

Unlike in the insulin targets, significantly enriched GO terms in the 20E-induced genes describe a variety of developmental processes (Fig. 4). Genes with these GO terms include components and targets of the Notch, Wg, EGF, and JAK/STAT pathways (Fig. 5). Also significantly affected was *sugarless*, which is required to synthesize the heparan sulfate moieties of proteoglycans needed for Dpp, Hh, FGF and Wg signaling (Hacker et al. 1997; Bornemann et al. 2004; Yan & Lin 2009). 20E, and not insulin, also supports the expression of a wide variety of developmentally important transcription factors, chromatin modifying proteins, and many genes that regulate tissue morphogenesis, including junctional, cytoskeletal and ECM components (Fig. 5). A smaller set of developmentally important genes seems to be regulated by either hormone; however 20E generally causes a more potent and long-lasting response in this group (Fig. 4 – Figure Supplement 1C). Of the genes that only 20E induces, roughly 30% are already known to perturb wing development when mutated.

**Figure 5:**
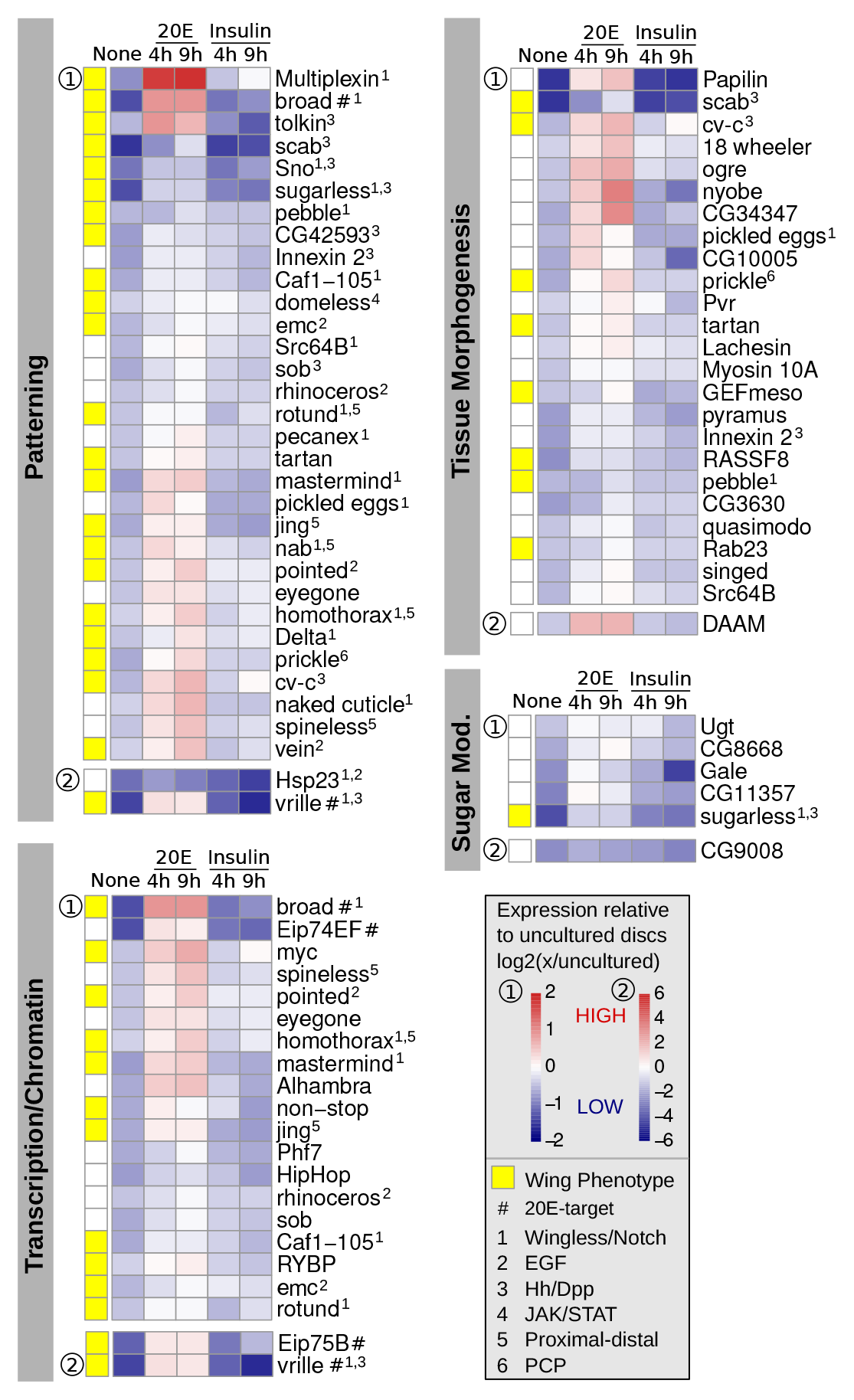
Patterning and morphogenesis genes that are regulated by 20E in culture. Normalized expression values are plotted across all conditions for selected 20E-regulated genes. The expression value of each gene (fkpm) is shown relative to its expression in uncultured discs, log2(x/uncultured). Genes were plotted on one of two scales (labeled as 1 or 2), depending on how much its values changed across conditions. Yellow squares indicate genes with published loss-of-function phenotypes in the wing. Previously identified 20E-targets and genes with known connections to specific patterning systems (in any context) are indicated.

On the whole, these transcriptional data suggest that insulin promotes proliferation through its well-understood effects on anabolic metabolism. However, 20E promotes proliferation over longer times while influencing only a subset of the direct growth promoting genes activated by insulin. Instead, 20E supports the expression of genes involved in wing patterning and morphogenesis.

### Perturbation of 20E signaling disrupts wing disc patterning *in vivo*

To determine whether our data from explant culture reflect an *in vivo* requirement for 20E in maintaining patterning systems during growth, we genetically perturbed 20E signaling *in vivo* and then assessed its effect on a broad range of morphogen and PCP pathways (Fig. 6). To do this, we expressed a dominant negative Ecdysone receptor (EcR-DN) in the dorsal compartment of wing imaginal discs for 24hr of the third larval instar. Consistent with previous results (Herboso et al. 2015), EcR-DN expression autonomously reduces tissue size and the number of mitotic cells (Fig. 6-Figure Supplement 1A). However, the tissue appears otherwise healthy – caspase staining reveals no elevation in apoptosis ((Herboso et al. 2015) and data not shown), and neither localization nor protein levels of Discs large is perturbed (Fig. 6A).

**Figure 6:**
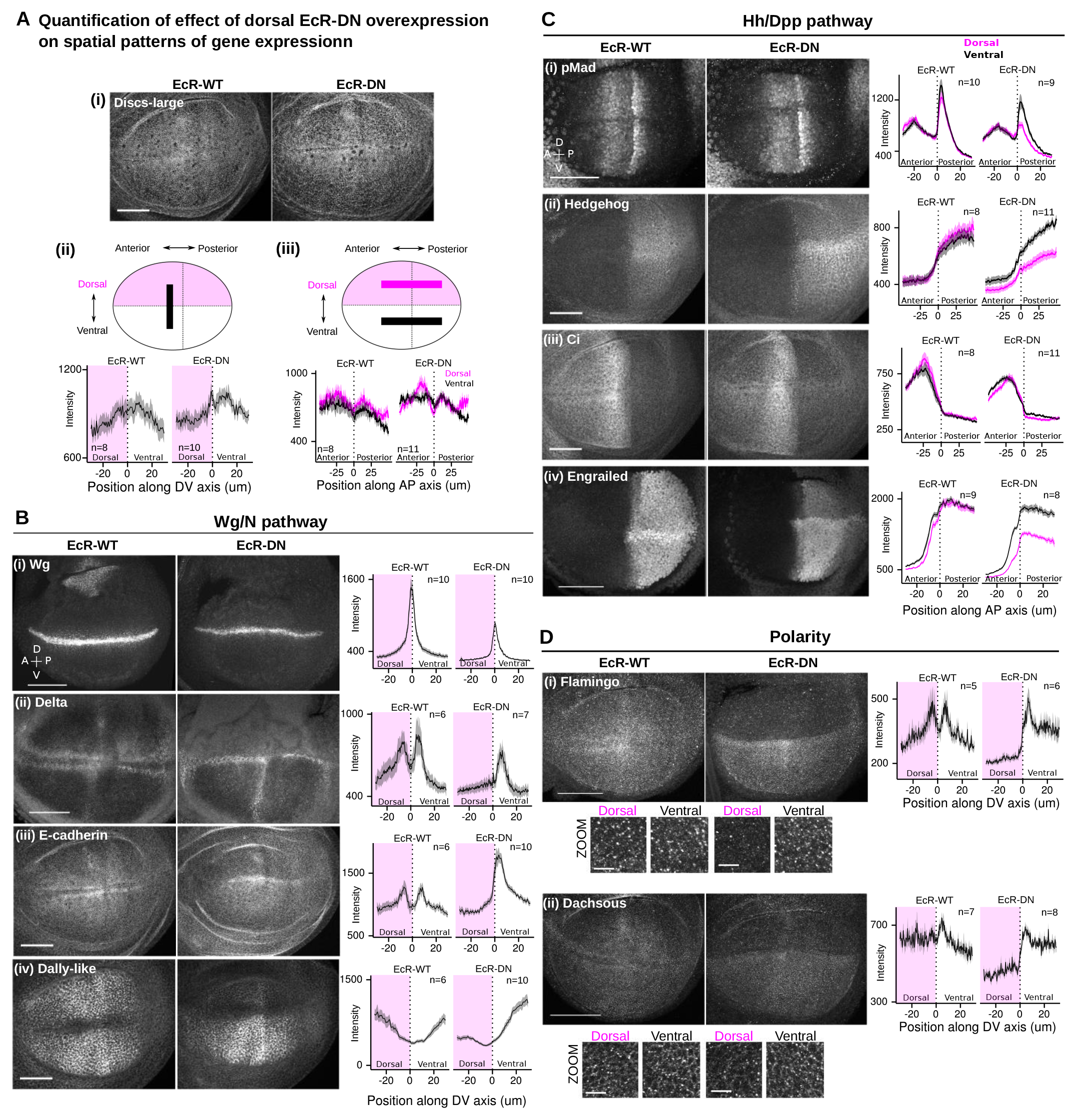
Widespread effects on morphogen signaling and PCP upon perturbation of 20E signaling *in vivo*. A dominant-negative allele of EcR-B1 (W650A, abbreviated EcR-DN) was overexpressed for 24hr in the dorsal compartment of the wing pouch using apterous-GAL4 and temperature-sensitive GAL80. As a negative control performed in parallel, we overexpressed the wild type EcR-B1 gene (EcR-WT), which does not significantly affect growth or proliferation of the disc (Fig. 6--Figure Supplement 1) (A). Quantification of the expression patterns in this experiment was done in one of two ways, each demonstrated in (A) for Discs-large (i), a septate junction marker. The first scheme (Aii) was used for proteins in (B) and (D): the intensity profile (arbitrary units) along a line in the Anterior compartment, drawn from Dorsal to Ventral, was compared between EcR-WT and EcR-DN expressing discs. For Discs-large (Aii), this profile is unaffected by EcR-DN expression. In contrast, intensity in the dorsal (pink shaded) compartments is altered for the morphogen Wg (Bi), the Notch ligand Delta (Bii), adhesion protein E-Cadherin (Biii), the HSPG Dallylike (Biv), the core PCP component Flamingo (Di) and the Fat PCP component Dachsous (Dii). The lower images of parts Di-ii are zoomed-in regions from the anterior compartment of the above images. The effect on the stripe of Wg expression was confirmed by expressing EcR-DN in the posterior compartment (Fig. 6--Figure Supplement 2). The second scheme (Aiii) was used for the Hh/Dpp pathways (C): the profile of absolute intensity along a line drawn from Anterior to Posterior in the Dorsal compartment (pink) was compared to that along a line drawn in the Ventral compartment (black). For Discs-large (Aiii), the black and pink lines are similar, both in EcR-WT (left) and EcR-DN (right) expressing discs. However, a readout of Dpp signaling, pMAD (Ci) was affected, particularly in the posterior compartment, by EcR-DN expression. In addition, compartment-autonomous differences in the Hh pathway were observed: at the level of Hh itself (Cii), its transcriptional coactivator Ci (Ciii), and the downstream target Engrailed (Civ). The effect of perturbing 20E signaling on Wg, Hh, Ci, Engrailed, and Discs-large were confirmed by an independent method (removal of circulating 20E) in Fig. 6--Figure Supplement 3. Scale bars for the zoomed regions in D are 20um; elsewhere they are 50um. In all quantified intensity profiles, the solid line represents the mean of several discs from the same experiment (precise # depicted in each graph), and the shaded ribbon is the standard deviation. All images are shown with dorsal up and anterior to the left.

Expressing EcR-DN in the dorsal compartment decreases Wg expression both in the hinge and in the stripe at the DV boundary (Fig. 6Bi). The autonomous effect of EcR-DN on the DV boundary Wg stripe is more obvious when it is expressed only in the posterior compartment (Fig. 6-Figure Supplement 2). The effect of EcR-DN expression clearly reflects a true requirement for 20E, as wing discs deprived of 20E *in vivo* have severely reduced levels of Wg and also less growth (Fig. 6—Figure Supplements 1B and 3A). These results are consistent with previously published data showing that EcR-DN expression can block Wg expression (Mirth et al. 2009; Herboso et al. 2015).

In wild type discs, both E-Cadherin and Delta are upregulated by Wg signaling on either side of the DV boundary (Micchelli et al. 1997; Jaiswal et al. 2006; de Celis & Bray 1997). Dorsal EcR-DN expression prevents both from accumulating on the dorsal side of the DV boundary, consistent with reduced Wg signaling there (Fig. 6Bii-iii). Like at the DV boundary, EcR-DN expression also affects elevated Delta expression on either side of a region of high Hedgehog signaling near the AP boundary (Fig. 6Bii). This effect on Delta is consistent with our transcriptomic data showing that mRNA levels of Delta itself, as well as effectors of the EGF and Notch pathways that regulate the AP Delta pattern, drop in culture without 20E (Fig 5).

Also consistent with our transcriptomic data, we found that Dally-like protein levels are considerably reduced in the EcR-DN expressing compartment (Fig. 6Biv). Given the broad effects of HSPG on morphogen signaling, we therefore also looked at Hh and Dpp pathway. Indeed, EcR-DN decreases levels of pMAD (a readout of Dpp signaling) in the posterior compartment (Fig. 6Ci). Furthermore, Hh protein levels and signaling are strongly affected (Fig. 6Cii-iv). Hh staining intensity decreases in the posterior producing cells, consistent with roles for Dally and Dally-like in controlling its trafficking there (Ayers et al. 2010; Ayers et al. 2012; Eugster et al. 2007). Furthermore, the gradient in the anterior receiving cells is diminished and the amplitude and range of Hedgehog signaling outputs are reduced. Ci is stabilized over a shorter distance from the AP boundary (Fig. 6Ciii). Engrailed expression is also reduced–both anterior to the AP boundary, where expression is activated by Hh signaling, and posteriorly where its expression is maintained independently of Hh by chromatin modifications (DeVido et al. 2008) (Fig. 6Civ). The Hh gradient and signaling output was severely reduced in wing discs from larvae deprived of 20E *in vivo*, confirming that the effects of EcR-DN expression reflect a real requirement for 20E (Fig. 6-Figure Supplement 3B-D).

Our transcriptomic analysis also revealed that 20E affects expression of the Core PCP component Prickle and the Fat PCP component Four-jointed (Fig. 5, Fig. 4-Figure Supplement 1). To examine global patterns of PCP upon EcR-DN expression, we stained discs with antibodies to Dachsous (Ds) and Flamingo (Fmi). Neither of these proteins was transcriptionally affected by 20E in culture, but their patterns of subcellular localization should reveal any disturbances in global PCP patterns. Strikingly, EcR-DN expression dramatically lowered protein levels of both Ds and Fmi, presumably post-transcriptionally, such that any residual planar polarity was unquantifiable (Fig. 6D).

In summary, both *in vivo* experiments and transcriptomic data from explanted discs suggest that 20E is required to maintain the signal transduction and planar cell polarity pathways that drive growth and patterning in the wing imaginal disc. Given its key role of regulating patterning *in vivo* and *in vitro*, addition of 20E to explant culture should be critical for measuring normal patterning of growth.

### Cellular contributions to oriented tissue growth

Having established that 20E supports longer proliferation and is better than insulin at maintaining the gene expression patterns underlying oriented growth, we used the new 20E-containing media to investigate the cell dynamics underlying growth and its anisotropy with live imaging. We analyzed three discs in 13hr timelapses.

We first examined area change in the wing pouch and its cellular contributions (Fig. 7Bi). We note that tissue area (blue line in Fig. 7Bi) decreases during the first two hours, almost entirely as a consequence of cell area changes (green line in Fig. 7Bi), suggesting that some time is required for discs to adapt to culture or recover from mounting. However, after the adaptation phase, tissue area grows. Cell divisions contribute positively to tissue area growth and are only partly counteracted by modest levels of cell extrusion (cyan line in Fig. 7Bi) and slight cell area reduction (Fig. 7Bi). This slight reduction in cell area is consistent with *in vivo* observations of growth over development time (Aegerter-Wilmsen et al. 2012; Mao et al. 2013).

**Figure 7:**
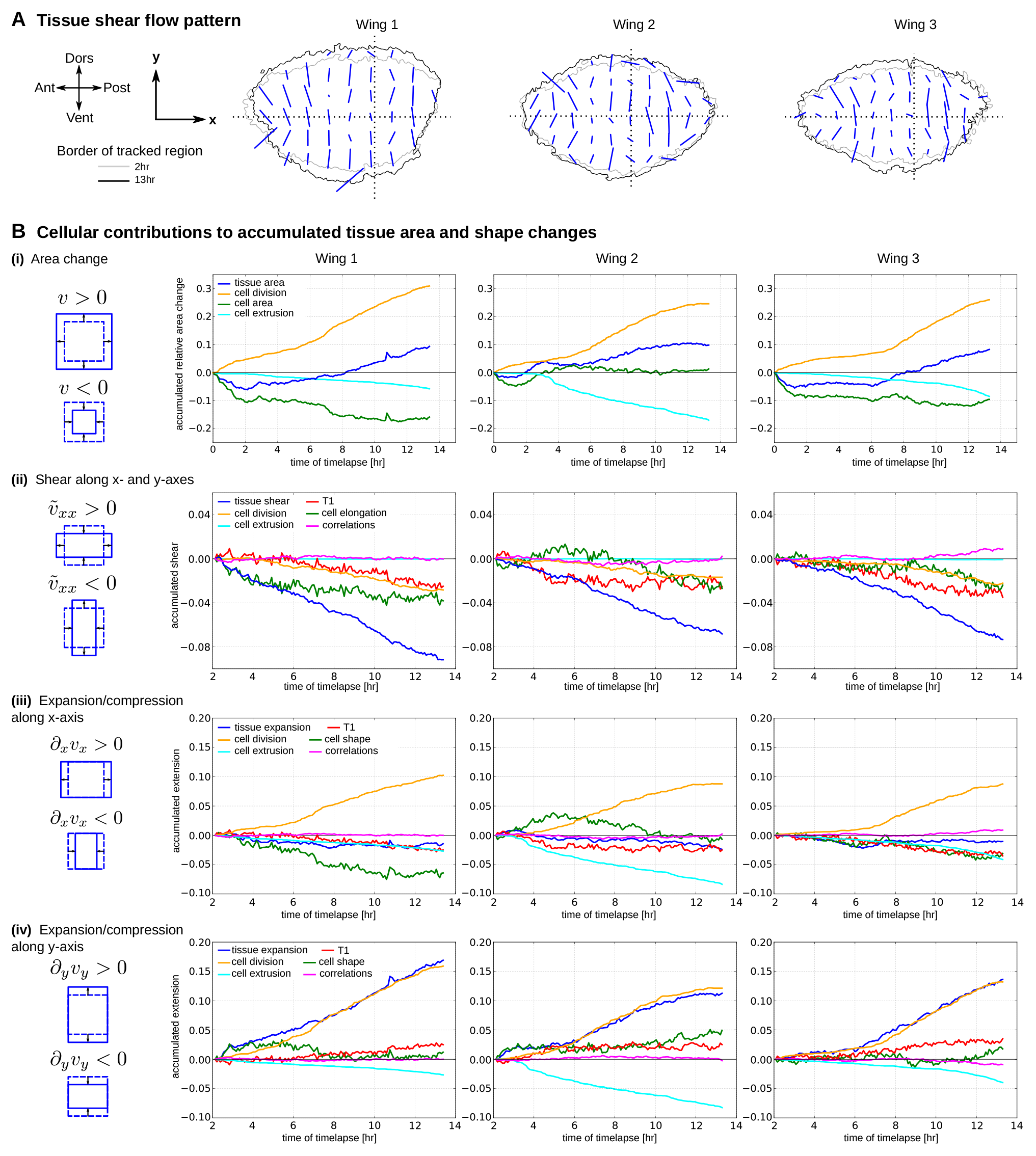
Quantification of tissue growth and its cellular contributions. Three E-Cadherin-GFP-expressing wing discs were imaged for >13hr, and cells in the future wing blade region were segmented and tracked. (A) Tissue shear was accumulated between 2hr and 13hr in local regions of the tissue defined by a grid. Accumulated shear is represented by blue bars plotted on outlines of the tracked region at 2hr (grey) and 13hr (black). (B) The trace of the velocity gradient tensor *υ* = *∂_x_υ_x_* + *∂_y_υ_y_* quantifies the relative change of tissue area. The diagonal component of the traceless symmetric part of the velocity gradient tensor 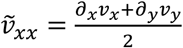 quantifies the tissue shear flow along x- and y-axes. (i) Relative change of tissue area, accumulated from the start of the timelapse, contains contributions from average cell area change, cell divisions and cell extrusions. In all experiments, positive tissue area growth (blue) is driven by cell divisions (orange) and partially opposed by cell extrusions (cyan) and cell area change (green). (ii) Accumulated tissue shear along the x-axis, averaged over the entire tracked region is plotted as a function of time (blue). The continuous shear of the blade perpendicular to the x-axis (negative values of blue curve) is composed of similar contributions from cell elongation change (green), T1 transitions (red) and cell divisions (orange). Contributions from cell extrusions (cyan) and correlations (magenta) are negligible. (iii-iv) Velocity gradient components, *∂_x_υ_x_* and *∂_y_υ_y_* represent tissue extension or compression along the x- and y-axis, respectively. We calculated the accumulated extensions along x- and y-axes from the relative area change of the tissue and the tissue shear. Cellular contributions to the tissue expansion along the x-axis were determined as described in the Methods. We find that the tissue does not extend along the x axis (blue line in iii), although cell divisions do contribute to tissue expansion along x axis (orange line in iii). In contrast, tissue extension along y axis (blue line in iv) closely follows the cell division contribution (orange line in iv). The sum of other contributions is small.

To analyze the anisotropy of tissue growth in 20E, we measured the patterns of pure tissue shear accumulated during 11hr after the 2hr adaptation phase using the Tissue Miner computational framework (Etournay et al. 2016) and Triangle Method (Merkel et al. 2017) described previously. Pure shear describes a change in aspect ratio irrespective of area change. In Fig. 7A, local cumulative tissue shear is indicated by a line whose length is proportional to the magnitude of pure shear and whose orientation indicates its axis. We observe that tissue shear is oriented perpendicular to an axis defined by the DV boundary throughout the tracked region of the wing pouch, generally consistent with analyses of *in vivo* clone data (González-Gaitán et al. 1994; Resino et al. 2002; Baena-López et al. 2005; Worley et al. 2013; Mao et al. 2013; Heemskerk et al. 2014).

To examine the dynamics of tissue shape change, and its underlying cellular contributions, we accumulated the average tissue shear and its cellular contributions over the entire tracked region using a coordinate system with an x-axis corresponding to the DV boundary (dashed horizontal line in Fig. 7A) and a y-axis perpendicular to it. In this coordinate system, a positive xx component of shear corresponds to shear parallel to the DV boundary, while a negative xx component of shear corresponds to shear perpendicular to the DV boundary. Fig. 7Bii (blue line) shows the average xx component of tissue shear throughout the tracked region, accumulated over time after the adaptation phase. These plots reveal that the wing pouch becomes more anisotropic as it grows. Examining the cellular contributions to tissue shear reveals that cell divisions (orange line in Fig. 7Bii) contribute to the change in shape of the wing pouch during growth. However, cell division orientation cannot entirely account for tissue shear: quantitatively similar contributions to tissue shear stem from both cell rearrangements (red line in Fig. 7Bii) and cell shape changes (green line in Fig. 7Bii).

Comparing the outlines of the wing pouch at the beginning and end of the movies (Fig. 7A) suggests that wing pouch tissue is expanding almost entirely perpendicular to the DV boundary. To separately quantify expansion along the x- and y-axes over time, we determined the xx and yy components of velocity gradient tensor based on the measured shear and area expansion (see Methods). Indeed, tissue expansion (blue lines in 7Biii and 7Biv) occurs along the y-axis, perpendicular to the DV boundary (Fig. 7Biv) and not along the x-axis (Fig. 7Biii). We decomposed the xx and yy components of the tissue velocity gradient tensor into their respective cellular contributions (see Methods). This calculation reveals that expansion of the wing pouch along the y-axis is almost identical to the expansion contributed by cell divisions in all three movies (compare orange and blue lines in Fig. 7Biv). Surprisingly, along the x-axis, the relationship between tissue flow and cell divisions is very different. Expansion due to cell divisions is only slightly smaller along the x-axis than along the y-axis (orange lines in Fig. 7Biii-iv). Nevertheless, the xx component of tissue velocity gradient is small and negative – i.e. the tissue does not expand in this axis but rather contracts slightly. In this direction, the sum of the other cellular contributions is larger than along the y-axis and balances the contribution from cell divisions. We note that although the contributions of cell shape changes, rearrangements and extrusions vary in each of the three movies, they always combine to cancel the expansion along the x-axis due to cell division.

In summary, we use the improved method for long-term cultivation of wing imaginal discs to directly measure the cellular contributions to tissue shape and area changes. This analysis revealed that contributions from cell divisions to tissue shape changes are not nearly as anisotropic as tissue flow and indicates that additional morphogenetic mechanisms exist to prevent expansion of the wing parallel to the DV boundary.

## DISCUSSION

This work considerably expands our understanding of the cell dynamics in the growing *Drosophila* wing disc, revealing important contributions from cell rearrangements and cell shape changes to oriented tissue growth. Our analysis provides the foundation for future experiments aimed at understanding how patterned gene expression and tissue mechanics regulate cellular behavior during tissue growth. Furthermore, our transcriptomic data from wing discs cultured in 20E or insulin provide a novel global view of the direct response of wing imaginal tissues to these hormones. These data have *in vivo* implications for the organismal coordination of growth and development and constitute a valuable resource for the planning of any type of experiment requiring wing disc culture.

### 20E supports normal growth of cultured wing discs, independently of insulin

Studying the cellular basis of tissue development requires prolonged, high time resolution imaging that is ideally entirely *in vivo*. However, the *Drosophila* wing grows during larval stages, when animals are mobile and feeding, and thus a completely *in vivo* approach to imaging growth is practically not feasible. While advances in clonal marking and imaging methods now allow the periodic monitoring of clone size and shape in the same animal over its development (Heemskerk et al. 2014), the available frame rate is still far too low to study cell dynamics. Analyzing whether cell boundary exchanges that are noisy and dynamic have an orientation, for example, demands the integration of high time resolution data over as much time as possible. Explant culture is the only option, provided that the patterning systems that drive oriented growth can be maintained.

While previous studies have used insulin to sustain discs long term in culture (Zartman et al. 2013; Mao et al. 2011; Mao et al. 2013; Legoff et al. 2013; Handke et al. 2014; Heller et al. 2016; Tsao et al. 2016), we show that the expression of numerous genes involved in patterning declines when discs are cultured without the steroid hormone 20E. 20E was previously thought to promote imaginal growth by repressing the expression of the translational inhibitor 4E-BP (Herboso et al. 2015); however, we show here that in addition, 20E broadly affects many different signaling pathways, each one of which is critical for growth. These data indicate that the inclusion of 20E in culture media should be critically important to the goal of replicating *in vivo* growth patterns. While high concentrations of 20E promote eversion, 10-fold lower concentrations promote growth in the absence of insulin. This condition actually supports proliferation for longer than insulin-containing media: up to ~24hrs in 20E, compared to only ~7-10hrs in insulin (Fig. 1, and (Zartman et al. 2013; Handke et al. 2014; Tsao et al. 2016)).

How does the growth of 20E-cultured discs compare to that *in vivo?* Although discs can proliferate in culture up to 24hr, they slow down sooner (after ~13hr) if they are continuously imaged. Nonetheless, for the first 13hr of live imaging, our direct quantification of the pattern of tissue growth agrees well with data generated using indirect methods to follow growth *in vivo* (González-Gaitán et al. 1994; Resino et al. 2002; Baena-López et al. 2005; Mao et al. 2013; Worley et al. 2013; Heemskerk et al. 2014). We also confirm the data from fixed tissues (Baena-López et al. 2005) showing that cell divisions are slightly biased to be perpendicular to the DV boundary, at least in central regions of the pouch (compare orange lines in Fig. 7Biii-iv). We estimate that the overall cell doubling time in continuously imaged explanted discs is about 3x longer than the estimated rate in well-fed animals (Garcia-Bellido & Merriam 1971; Bryant & Levinson 1985; González-Gaitán et al. 1994; Heemskerk et al. 2014). However, phototoxicity associated with continuous imaging seems to affect not just the duration but also the rate of proliferation: estimating the rate from the density of PH3-stained nuclei in discs that were not continuously imaged suggests a value that is closer to that of freshly explanted discs (Fig. 1).

Interestingly, *in vivo*, if animals are starved after the attainment of a critical weight, the release of insulin-like peptides from the brain is reduced (Géminard et al. 2009) and growth arrests in the larval – but not imaginal – tissues (Britton & Edgar 1998; Cheng et al. 2011). The rate of imaginal proliferation during this so-called “sparing” condition is ~60% lower than in the well-fed state (Cheng et al. 2011). Thus, proliferation of explants given 20E but not insulin may be closer to this *in vivo* low-insulin state. Importantly, while the ablation of insulin-producing cells in the brain reduces the size of the emerging adult wings, these wings are nevertheless well-proportioned (Rulifson et al. 2002). Thus, the orientation of tissue growth and its underlying cell dynamics is likely to be preserved in discs cultured with 20E in the absence of insulin. Perhaps the slight but reproducible lag in the division rate at early times of 20E culture (Fig. 1Biii and Fig. 7Bi) reflects a transition phase when discs taken from well-fed animals must switch to an insulin-independent/20E-dependent mode of growth.

### Implications for hormonal regulation of growth

The mechanism underlying the sparing of imaginal tissues is unknown. Sparing of the central nervous system involves a locally provided ligand that activates downstream signaling in the insulin pathway (Cheng et al. 2011). Our transcriptomic data provide no evidence that wing discs can produce their own insulin-like ligands or that they activate insulin signaling in the absence of exogenous ligands. Nonetheless, wing discs in culture continue to proliferate in 20E, even when no insulin-like ligand is present. This finding raises the possibility that 20E is also sufficient to support their proliferation *in vivo* during starvation. While 20E promotes imaginal growth, it inhibits that of larval tissues (Delanoue et al. 2010), potentially explaining the opposite response to starvation of these two tissue types.

Under well-fed conditions, insulin signaling must be able to combine with 20E signaling to further increase imaginal disc growth, but this feature is not reproduced in culture (Fig. 1Biv) and (Handke et al. 2014). *Drosophila* has 7 Insulin-like peptides that interact not only with the single receptor, but also with other proteins that modulate their activities (Honegger et al. 2008; Arquier et al. 2008; Okamoto et al. 2013). Perhaps specific insulin peptides or their binding proteins allow proper interfacing with 20E signaling to promote growth.

### Cellular dynamics underlying oriented tissue growth in the wing disc

While the mechanisms dictating cell division orientation in this tissue have been studied (Mao et al. 2011; Mao et al. 2013; Legoff et al. 2013), it has remained unclear to what extent the observed slight bias in cell division orientation could account for the orientation of tissue growth. Is cell division the only behavior that contributes to oriented tissue growth? The Triangle Method (Merkel et al. 2017; Etournay et al. 2015) for decomposing tissue shape changes into quantitative contributions from all types of cellular events provides the measurement tool to connect the cell and tissue scales. Our results reveal that the observed bias in cell division orientation in the central region of the disc is insufficient to explain the anisotropy of tissue growth. While expansion of the tissue perpendicular to the DV axis is driven primarily by cell divisions, in the orthogonal direction, cell shape changes and T1 transitions fully cancel the contribution to expansion by cell divisions, such that the wing does not expand along an axis parallel to the DV boundary.

Previous efforts to quantify cell rearrangements have used live imaging of discs cultured in insulin and have concluded that they occur either with very low frequency (Gibson et al. 2006; Legoff et al. 2013), or with a significant frequency but no pattern or orientation at the tissue-scale (Mao et al. 2013; Heller et al. 2016). In contrast, we see that the contribution of cell rearrangements to tissue growth is quantitatively similar to that of oriented cell division and helps restrict growth parallel to the DV boundary. This discrepancy could either come from the differing culture conditions (lack or presence of 20E) or to the inherent difficulties in defining and counting cell rearrangements. A classic T1 transition occurs when two pairs of cells exchange neighborships: the connecting bond between two cells shrinks as all cells come together to make a 4-way vertex; then, a new bond is formed such that the two cells that were originally separated come into contact, while the two that were originally together stay separate. But sometimes the 4-way vertex simply resolves back without changing the original neighborship, and sometimes the neighborship fluctuates back and forth several times. How and when do you count the rearrangement? The method that we use overcomes the problem of having to classify a “true” T1 event based on observation time windows. It considers all changes in neighborship, regardless of the length of time they persist, and quantifies their accumulated contribution to tissue shape change. Note that the shear decomposition method we use does not simply quantify the number and orientation of T1s but the tissue shape change that they cause. This approach may be key to our ability to detect and quantify the important contribution of cell rearrangements to oriented growth of the wing disc.

What mechanisms might allow cell rearrangements, cell shape changes and cell extrusions to cancel expansion due to cell division specifically in one direction? We propose that a mechanical constraint prevents wing pouch expansion parallel, but not perpendicular, to the DV boundary. In this case, area growth due to cell division would lead to anisotropic expansion with corresponding anisotropic stresses that could be associated with oriented cell shape changes and rearrangements. Consistent with this idea, we observe that the sum of expansion due to rearrangements, shape changes and extrusions is more reproducible than their individual cellular contributions. Long range anisotropic stresses could coordinate these cellular processes and account for their mutual dependence. A similar dependence between cellular contributions is observed in the pupal wing, where the constant tissue area is maintained by epithelial tension via extracellular matrix connections to the cuticle (Etournay et al. 2015). We do not yet know whether the larval wing disc is mechanically constrained. It will be interesting to explore whether the extracellular matrix has anisotropic properties or whether the folding pattern of the tissue influences growth orientation.

The fact that Fat PCP mutant wings are associated with less anisotropic clone shapes and more weakly oriented cell divisions (Baena-López et al. 2005; Mao et al. 2011) has led to the idea that Fat orients growth through its effect on cell division orientation. We find that shear due to cell divisions cannot fully account for the anisotropy of growth, however. This result suggests either that there are other orienting factors, or that Fat PCP has more profound effects on growth anisotropy than can be accounted for by cell division.

Overall, this work open new avenues towards an understanding of wing morphogenesis that integrates different scales – from the larval hormonal networks, to the morphogen-dependent patterns of cell dynamics from which tissue shape emerges.

### Figure legends

**Figure 1: 20E is sufficient to stimulate prolonged cell division in cultured wing discs**

Wing discs from mid-third instar larvae (96hr AEL) were cultured for indicated times and then stained with phospho-Histone H3 to detect mitotic nuclei at the indicated timepoints. (A) Representative images and (B) their quantification. Scale bar indicates 50μm. Discs were given (i) no hormone supplement, (ii) 5μg/ml insulin, (iii) 20nM 20E or (iv) both. In (v) is an image of an uncultured disc from the beginning of the experiment (96hr AEL). In (B), we show pooled results from multiple experiments from different days. Each dot is the mean density of mitotic nuclei (# nuclei per 10x10μm^2^ area, averaged across the pouch from a single disc). Discs from experiments performed on the same day have the same color. The box plots summarize all data for each condition. Hinges correspond to the first and third quartiles; lines extend to the highest and lowest values within 1.5*IQR (inter-quartile range); outlier points are black. n.d. = not determined. (C) Apical cell area the wing pouch after 9hr of culture. Overall, the morphology is maintained during culture, except in presence of both hormones. Lowering the concentration of 20E in combination with insulin does not further prolong proliferation (See Fig. 1--Figure Supplement 1).

**Figure 2: Size and overlap of the direct transcriptional responses to 20E or insulin**

Transcriptomes of wing discs cultured in either 20E or insulin were sequenced and compared to those of uncultured discs and discs cultured without any hormone. A gene was classified as hormone-dependent if its levels changed upon culture without hormone and responded in the correct direction (more similar to uncultured levels) when cultured with hormone. (A) Total number of genes that responded to 20E or insulin at two timepoints. (B) Overlap between the hormone-dependent responses at 4hr (left), 9hr (right) or when each hormone supports the most amount of proliferation (4hr for insulin, 9hr for 20E). Circles are all drawn to scale. Numbers denote the amount of genes in the overlapping or non-overlapping sets.

**Figure 3: Processes that are transcriptionally regulated by insulin in cultured discs**

Shown are the Biological Process (BP) Gene Ontology (GO) terms that are significantly enriched in the set of genes identified to be activated (A) or repressed (B) by insulin in cultured wing discs (simplified with a semantic similarity algorithm; threshold=0.95). In both (A) and (B), we plot the GO terms according to the fraction of its total genes that were identified in the data (# genes in the data with this GO term/total size of the GO term). The size of the point indicates the significance of enrichment (BH-corrected *p*-value; cutoff of *p* < 0.005). The color indicates either 4hr (light orange in A, green in B) or 9hr of culture (dark orange in A, dark green in B). Note that by 9hr of culture in insulin, no GO terms are enriched in the group of genes repressed by insulin (B). Asterisks indicate terms shared by both insulin and 20E. Note that many categories are only enriched (or more significantly enriched) at 4hr. This transience in the response to insulin must result from downregulation of the pathway at a point downstream of PI3K activation, since a fluorescent reporter of this enzyme’s activity indicates that it is still active for the duration of culture (See Fig. 3--Figure Supplement 2).

**Figure 4: Processes that are transcriptionally regulated by 20E in cultured discs**

Shown are the Biological Process (BP) Gene Ontology (GO) terms that are significantly enriched in the set of genes identified to be activated (A) or repressed (B) by 20E in cultured wing discs (simplified with a semantic similarity algorithm; threshold=0.9). In both (A) and (B), we plot the GO terms according to the fraction of its total genes that were identified in the data (# genes in the data with this GO term/total size of the GO term). The size of the point indicates the significance of enrichment (BH-corrected *p*-value; cutoff of *p* <0.005). The color indicates either 4hr (light orange in A, green in B) or 9hr of culture (dark orange in A, dark green in B). Asterisks indicate terms shared by both insulin and 20E.

**Figure 5: Patterning and morphogenesis genes that are regulated by 20E in culture**

Normalized expression values are plotted across all conditions for selected 20E-regulated genes. The expression value of each gene (fkpm) is shown relative to its expression in uncultured discs, log_2_(x/uncultured). Genes were plotted on one of two scales (labeled as 1 or 2), depending on how much its values changed across conditions. Yellow squares indicate genes with published loss-of-function phenotypes in the wing. Previously identified 20E-targets and genes with known connections to specific patterning systems (in any context) are indicated.

**Figure 6: Widespread effects on morphogen signaling and PCP upon perturbation of 20E signaling *in vivo***

A dominant-negative allele of EcR-B1 (W650A, abbreviated EcR-DN) was overexpressed for 24hr in the dorsal compartment of the wing pouch using apterous-GAL4 and temperature-sensitive GAL80. As a negative control performed in parallel, we overexpressed the wild type EcR-B1 gene (EcR-WT), which does not significantly affect growth or proliferation of the disc (Fig. 6--Figure Supplement 1) (A). Quantification of the expression patterns in this experiment was done in one of two ways, each demonstrated in (A) for Discs-large (i), a septate junction marker. The first scheme (Aii) was used for proteins in (B) and (D): the intensity profile (arbitrary units) along a line in the Anterior compartment, drawn from Dorsal to Ventral, was compared between EcR-WT and EcR-DN expressing discs. For Discs-large (Aii), this profile is unaffected by EcR-DN expression. In contrast, intensity in the dorsal (pink shaded) compartments is altered for the morphogen Wg (Bi), the Notch ligand Delta (Bii), adhesion protein E-Cadherin (Biii), the HSPG Dallylike (Biv), the core PCP component Flamingo (Di) and the Fat PCP component Dachsous (Dii). The lower images of parts Di-ii are zoomed-in regions from the anterior compartment of the above images. The effect on the stripe of Wg expression was confirmed by expressing EcR-DN in the posterior compartment (Fig. 6--Figure Supplement 2). The second scheme (Aiii) was used for the Hh/Dpp pathways (C): the profile of absolute intensity along a line drawn from Anterior to Posterior in the Dorsal compartment (pink) was compared to that along a line drawn in the Ventral compartment (black). For Discs-large (Aiii), the black and pink lines are similar, both in EcR-WT (left) and EcR-DN (right) expressing discs. However, a readout of Dpp signaling, pMAD (Ci) was affected, particularly in the posterior compartment, by EcR-DN expression. In addition, compartment-autonomous differences in the Hh pathway were observed: at the level of Hh itself (Cii), its transcriptional coactivator Ci (Ciii), and the downstream target Engrailed (Civ). The effect of perturbing 20E signaling on Wg, Hh, Ci, Engrailed, and Discs-large were confirmed by an independent method (removal of circulating 20E) in Fig. 6--Figure Supplement 3. Scale bars for the zoomed regions in (D)are 20um; elsewhere they are 50um. In all quantified intensity profiles, the solid line represents the mean of several discs from the same experiment (precise # depicted in each graph), and the shaded ribbon is the standard deviation. All images are shown with dorsal up and anterior to the left.

**Figure 7: Quantification of tissue growth and its cellular contributions**

Three E-Cadherin-GFP-expressing wing discs were imaged for >13hr, and cells in the future wing blade region were segmented and tracked. (A) Tissue shear was accumulated between 2hr and 13hr in local regions of the tissue defined by a grid. Accumulated shear is represented by blue bars plotted on outlines of the tracked region at 2hr (grey) and 13hr (black). (B) The trace of the velocity gradient tensor *v* = *∂_x_υ_x_* + *∂_y_υ_y_* quantifies the relative change of tissue area. The diagonal component of the traceless symmetric part of the velocity gradient tensor 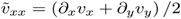 quantifies the tissue shear flow along x-and y-axes. (i) Relative change of tissue area, accumulated from the start of the timelapse, contains contributions from average cell area change, cell divisions and cell extrusions. In all experiments, positive tissue area growth (blue) is driven by cell divisions (orange) and partially opposed by cell extrusions (cyan) and cell area change (green). (ii) Accumulated tissue shear along the x-axis, averaged over the entire tracked region is plotted as a function of time (blue). The continuous shear of the blade perpendicular to the x-axis (negative values of blue curve) is composed of similar contributions from cell elongation change (green), T1 transitions (red) and cell divisions (orange). Contributions from cell extrusions (cyan) and correlations (magenta) are negligible. (iii-iv) Velocity gradient components *∂_x_υ_x_*, and *∂_y_υ_y_* represent tissue extension or compression along the x- and y-axis, respectively. We calculated the accumulated extensions along x- and y-axes from the relative area change of the tissue and the tissue shear. Cellular contributions to the tissue expansion along the x-axis were determined as described in the Methods. We find that the tissue does not extend along the x axis (blue line in iii), although cell divisions do contribute to tissue expansion along the x axis (orange line in iii). In contrast, tissue extension along the y axis (blue line in iv) closely follows the cell division contribution (orange line in iv). The sum of other contributions is small.

## MATERIALS AND METHODS

### Flies

The culture experiments described in Fig. 1 and the transcriptomic analyses were performed using wild type OregonR *Drosophila*. For live imaging experiments, we used animals in which the endogenous E-Cadherin gene had been fused with GFP (Huang et al. 2009). To enable temperature inducible GAL4-dependent gene expression, we combined the *apterous-GAL4* (Marois et al. 2006), *phantom-GAL4* (Ono et al. 2006) and *engrailed-GAL4* (DeVido et al. 2008) loci with the temperature-sensitive GAL4 repressor GAL80^ts^ driven by the tubulin promoter. UAS-EcR.B1 (BDSC 6469) and UAS-EcR.B1-DeltaC655.W650A (BDSC 6872) were acquired from Bloomington Stock Center (Bloomington, Indiana, USA). The line used to induce RNAi against *neverland* was obtained from the group of R. Niwa (Yoshiyama et al. 2006).

Flies were grown on standard media containing cornmeal, molasses agar and yeast extract under a 12hr light/dark cycle at 25°C (except when gene expression was induced using the temperature-sensitive GAL80, described below).

For all disc culture experiments, we used wing discs from larvae at 96hr after egg laying (AEL). Flies were allowed to lay eggs on apple juice agar plates (supplemented with yeast paste) for 1-2hr at 25°C. Agar pieces containing ~10-12 eggs each were placed into food vials and grown at 25C°C. The middle of the egg collection window was considered to be 0hr AEL.

### Wing disc culture

#### Culture media

Early experiments (data not shown) indicated that overall disc morphology was best preserved in Grace’s Insect culture medium, compared to Schneider’s or Shield’s and Sang. Thus, Grace’s was used for all experiments. Grace’s medium (Sigma G9771) was prepared without Sodium bicarbonate but with the addition of 5mM BisTris. The pH was adjusted to 6.6-6.7 at room temperature, and the prepared liquid media was stored at 4°C for no longer than one month. On the day of the experiment, we added 5% Fetal Bovine Serum (ThermoFisher/Invitrogen 10270098) and Penicillin-Streptomycin (Sigma P4333, 100x) to impede microbial growth. Note that while other groups have included female fly extract in the media (Zartman et al. 2013; Handke et al. 2014; Tsao et al. 2016), we found it to be less reproducible than FBS (data not shown). Furthermore, recent evidence indicates that fly extract is associated with non-physiological calcium waves in explanted wing discs (Balaji et al. 2017).

Stock solutions of either 20E or insulin were added just prior to the start of the experiment. 20E (Sigma H5142) was added as a 1:1000 dilution from an ethanol stock solution (stored at −20°C, prepared fresh weekly) to give a final concentration of 20nM in the media (except when otherwise noted). Bovine insulin (Sigma I5500) was prepared as a 10mg/ml stock solution in acidified water (stored at −20°C) and added at a final concentration of 5μg/ml. This concentration of insulin is comparable to the levels used in previous studies aimed at prolonging proliferation of discs in culture (Zartman et al. 2013; Handke et al. 2014). Given that our data on the proliferation index in discs cultured in insulin alone (Fig 1Bii) quantitatively agree with the results of these previous studies, the other differences between our culture media and theirs do not seem to be as important as hormonal content.

#### Culture procedure

To prepare larvae for dissection, we first floated them out of the food by adding 30% sucrose. Larvae were transferred using a brush or wide-tip transfer pipet into glass dishes. Excess food was removed by washing in sucrose, followed by distilled water. They were then surface sterilized by immersion in 70% ethanol for 1-2 min. Finally, they were washed once more with sterile water and then media. Except for the live imaging experiments, discs were dissected in hormone-free media and then immediately transferred from the dissecting well into media containing hormone. For each experiment, dissections were performed over a period of no more than 45min, and the start time of the culture was considered to be the midpoint of that dissection window. The number of discs analyzed per sample was limited by how many could be manually dissected as carefully as possible during that time window.

Discs were cultured in glass wells (Electron Microscopy Sciences, Hatfield, PA; 70543-30) in 500μl of growth media, with 10-20 discs per well, at 25°C in humidified chambers. Media was exchanged approximately every 2hr, except during the long incubations of 16-24hr, when there was a maximum incubation without media change of 8hr. At the end of culture, discs were fixed by exchanging the media with 4% PFA (8% PFA stock solution in PBS diluted 2X into Grace’s media). Discs were incubated for 20min at room temperature and stained as described below in the immunofluorescence section.

The data presented in Fig. 1 represent a compilation of results from multiple days (labeled in different colors). Two replicates were done for each hormonal condition and timepoint. All discs were included in the analysis, except in rare cases when a disc was distorted during mounting.

### Transcriptomic analysis

#### Sample preparation

Three biological replicates were performed for each hormone condition and timepoint. For each, ~20-30 discs were dissected and cultured as described above and then transferred to a microfuge tube in a volume of 15-20μl (growth media). Discs were frozen immediately in liquid nitrogen and stored at −80°C until all samples were ready for RNA isolation. A Qiagen RNeasy Mini kit was used to isolate RNA for sequencing. Samples were lysed by adding Buffer RLT + 2-mercaptoethanol and vortexing for 20-30 seconds on ice. The manufacturer’s instructions were followed for the rest of the protocol. Further purification was achieved with an ethanol precipitation.

#### Sequencing

mRNA was isolated from 1μg total RNA by poly-dT enrichment using the NEBNext Poly(A) mRNA Magnetic Isolation Module according to the manufacturer’s instructions (New England Biolabs). Final elution was done in 15μl of 2x first strand cDNA synthesis buffer. After chemical fragmentation by incubating for 15 min at 94°C, the sample was directly subjected to the workflow for strand-specific RNA-Seq library preparation (Ultra Directional RNA Library Prep, New England Biolabs). For ligation, custom adaptors were used (Adaptor-Oligo 1: 5′-ACA-CTC-TTT-CCC-TAC-ACG-ACG-CTC-TTC-CGA-TCT-3′, Adaptor-Oligo 2: 5′-P-GAT-CGG-AAG-AGC-ACA-CGT-CTG-AAC-TCC-AGT-CAC-3′). After ligation, adapters were depleted with an XP bead purification (Beckman Coulter), adding beads in a ratio of 1:1. Indexing was done during the following PCR enrichment (15 cycles) using custom amplification primers carrying the index sequence indicated with ‘NNNNNN′. (Primer1: Oligo_Seq 5′-AATGATACGGCGACCACCGAGATCTACACTCTTTCCCTACACGACGCTCTTCCGATCT-3′, primer2: 5′-GTGACTGGAGTTCAGACGTGTGCTCTTCCGATCT-3′, primer3: 5′-CAAGCAGAAGACGGCATACGAGAT NNNNNN GTGACTGGAGTT-3′. After two more XP beads purifications (1:1), libraries were quantified using Qubit dsDNA HS Assay Kit (Invitrogen). Equimolar amounts of each sample were pooled and distributed amongst four lanes of an Illumina HiSeq 2500 sequencer for 75bp single-end sequencing. On average we achieved 37 million reads per sample.

#### Processing of sequencing data

Short read data was trimmed using Cutadapt (Martin 2011) and aligned using TopHat2 (Kim et al. 2013). Differential gene expression analysis was performed for all pairwise comparisons between conditions and timepoints using Cufflinks (Trapnell et al. 2012). Our threshold for significance was q<0.01. We considered a gene to be expressed if it had a normalized count of fpkm (fragments per kilobase of transcript per million mapped reads) > 5.

Raw sequencing data and fpkm normalized expression data files have been deposited in NCBI’s Gene Expression Omnibus and are accessible through GEO Series accession number GSE92933.

#### Analysis of hormone-responsive genesets

The identification of hormone-responsive genesets was achieved by selectively filtering the differentially expressed genes table from Cufflinks using custom R scripts. First, we identified the genes in this table whose levels were significantly different (q<0.01) between no culture (freshly dissected discs) and culture without hormone. We split this culture-responsive subset into two groups based on whether their levels in uncultured discs were abnormally low or high after culture without hormone. To identify hormone-activated genes, we first identified the genes whose expression was abnormally low in culture with no hormone (relative to uncultured control). Amongst these, we identified genes whose levels were significantly elevated (either at 4hr or 9hr) when discs were culture with hormone (compared to culture with no hormone). Hormone-repressed genes had the opposite pattern: abnormally high when cultured without hormone but significantly lower in culture with hormone.

The Venn diagrams presented in Fig. 2 were generated using the VennDiagram package of R (Chen & Boutros 2011) after identifying the genes that were unique or common in each pairwise comparison.

We identified enriched Gene Ontology (GO) terms (Biological Process and Cellular Component) within each hormone-responsive dataset using the R-package ClusterProfiler (v 1.9) (Yu et al. 2012). To reduce redundancy, we used the simplify function of this package; similarity and p-value cutoffs are written in the figure legends. We also required enriched GO terms to have > 5 genes included in the data.

For selected genes, we plotted the expression levels across all conditions (Fig. 3-Figure Supplement 1, Fig. 4-Figure Supplement 1, Fig. 5) by taking the log2 of the normalized counts (fkpm) in each condition divided by that in the uncultured wing discs.

For the 589 genes regulated by 20E (either exclusively or also by insulin), we manually classified functional groupings after reviewing related literature, GO term assignments, and Flybase annotations. The classifications used in Fig. 4-Figure Supplement 1 and Fig. 5 are explained as follows: DNA replication/cell cycle includes anything having a positive effect on DNA replication and/or cell cycle progression. Cell death/DNA damage includes genes annotated with a function in apoptosis, programmed cell death, or DNA repair. Protein production includes splicing, translation, ribosome biogenesis, and protein folding. Nutrient transport was limited to proven or predicted members of the SLC/MFS superfamily. The patterning group includes anything known to regulate or be a target of developmental signaling pathways (in any system, not limited to the wing disc). The transcription/chromatin class included verified and putative transcription factors and chromatin modifying proteins. Tissue morphogenesis includes genes involved in cell-cell or cell-ECM contact, as well as cytoskeletal components and regulators. Lastly, the sugar modifications group describes anything involved in protein glycosylation. In Fig. 4-Figure Supplement 1, TOR targets were identified in (Teleman et al. 2008; Guertin et al. 2006), and FOXO targets were from (Teleman et al. 2008; Puig et al. 2003; Spellberg et al. 2015; Alic et al. 2014). In Fig. 5, 20E targets are based on (Beckstead et al. 2005; Gauhar et al. 2009).

After examining these expression profiles, we realized that there were 11 genes that appeared to change over time irrespective of hormone content and were thus falsely identified as being 20E-responsive. We removed these genes from consideration in the heatmaps by identifying those that responded equally well to both insulin and 20E at 9hr but to neither at 4hr.

### Temperature-sensitive GAL4/GAL80^ts^ experiments

*Apterous-GAL4, GAL80^ts^/CyO-GFP* females were crossed with males of *UAS-EcR* or *UAS-EcR-W650A*, and their progeny were kept at 18°C (permissive for GAL80 repressor function) on bromophenol blue-containing food for 7-8 days. Vials were then moved to 29°C (restrictive) to induce GAL4-dependent transcript of EcR constructs for 24hr. Upcrawling larvae were selected for dissection based on the absence of GFP and the presence of blue food in the gut (indicating that they are still many hours from pupariation) (Andres & Thummel 1994).

To remove circulating 20E, *UAS-CD8-GFP;;phantom-GAL4, GAL80^ts^/TM3-GFP* females were crossed with males of the *neverland* targeting line, and their progeny were kept at 18°C. Vials were shifted to 29°C, and discs from upcrawling larvae lacking the *TM3-GFP* balancer were dissected 2 or 4 days later. As controls, we used progeny of *UAS-CD8-GFP;;phantom-GAL4, GAL80^ts^/TM3-GFP* females crossed to *w^1118^* flies in parallel. The progeny of the control cross all pupariated by 2 days after shift to 29°C.

### Immunofluorescence

Primary antibodies used were: anti-Ci-full length (1:100, rat, DSHB 2A1 conc), anti-Dally-like (1:100, mouse, DSHB 13G8 conc), anti-Dachsous (1:1000, mouse, (Merkel et al. 2014)), anti-Delta (1:800, mouse, DSHB C594.9B, conc), anti-Engrailed (1:100, mouse, DSHB 4D9 conc), anti-Discs-large (1:100, mouse, DSHB 4F3), anti-E-Cadherin (1:100, rat, DSHB DCAD2 sup), anti-Flamingo (1:200, rabbit, (Sagner et al. 2012)), anti-Hh (1:500, rabbit, (Eugster et al. 2007)), anti-Patched (1:100, mouse, DSHB Apa1 conc), anti-pHistoneH3-Ser10 (1:1000, rabbit, Cell Signaling, Danvers, MA, USA), anti-pMAD (1:1000, rabbit, Epitomics, Burlingame, CA, USA), anti-Wg (1:100, mouse, DSHB 4D4). DSHB stands for Developmental Studies Hybridoma Bank (Iowa, USA).

Secondary antibodies used were: Alexa-fluor-488 goat anti-Rabbit (1:1000, ThermoFisher A11034), Alexa-fluor-555 goat anti-Mouse (1:1000, ThermoFisher A21424), Alexa-fluor-647 goat anti-Rat (1:500, ThermoFisher A21247)

Discs were dissected in PBS and then fixed for 20min at room temperature in 4% PFA/PBS. Samples were permeabilized by rinsing twice with PBS+0.1% TritonX100 (TPBS) and blocked with TPBS +0.1mg/ml BSA + 250mM NaCl for 45-60min at room temperature. They were then incubated overnight at 4°C in primary antibody solution made in PBS+0.1% TritonX100+0.1mg/ml BSA (BBX). The next day, stainings were washed in BBX, followed by BBX+4% normal goat serum. Samples were then incubated at room temperature for 2-4hr in the dark with secondary antibodies diluted in BBX. Finally, samples were washed several times in TPBS and mounted between a glass slide and coverglass separated by a double-sided tape spacer. The discs were oriented with their apical surface toward the coverglass and immersed in VectaShield mounting medium (Vector Laboratories H-1000).

Imaging of fixed samples was performed using an Olympus FV1000 laser scanning confocal microscope fitted with an Olympus BX61 inverted stand and motorized XYZ stage, driven by FV10-ASW 1.7 software. Wing discs were imaged using either an Olympus UApochromat 40X 1.35NA oil immersion or Olympus UPlanSApochromat 60X 1.35NA oil immersion objective.

All discs were imaged and analyzed, except in very rare cases when the discs were damaged or misshapen during mounting.

### Analysis of immunofluorescence images

#### Phospho-Histone H3

Z-stacks of wing discs stained with PH3, E-Cadherin, Patched and Wingless were analyzed in 2D after a maximum intensity projection. PH3+ nuclei were segmented using FIJI Weka segmentation plugin (Arganda-Carreras et al. 2017). A training set containing images from each condition of the experiment was used to generate a common classifier, which was then applied to all images. Probability masks were thresholded by likelihood and size. We quantified signal only in the wing pouch, which was identified morphologically by the innermost folds in the epithelium. Compartment and wing pouch size in Fig. 6-Figure Supplement 1 was measured on a 2D projection, using FIJI to manually identify the region of interest. Plotting was performed in R.

#### Cell area in fixed samples

In Fig. 1C, apical cell areas were determined and plotted using Tissue Analyzer (Aigouy et al. 2016) to segment cells based on E-Cadherin staining, and Tissue Miner (Etournay et al. 2016) to determine the area and position of each cell.

#### Intensity profiles

The changes to patterns of protein production/localization upon *in vivo* perturbation of 20E signaling were analyzed in a 2D maximum intensity projectio.n Using Fiji (Schindelin et al. 2012), a line 50 pixels wide (~20μm) was drawn either from dorsal to ventral in the anterior and posterior compartments or from anterior to posterior in the dorsal and ventral compartments. Absolute intensity values along these lines were averaged across all samples (imaged on the same day, under the same acquisition settings). Plotted is the mean and standard deviation of intensity along these lines. At least five discs were imaged per biological condition; exact numbers are shown in the figures.

### Long-term timelapse imaging

Discs were dissected directly in growth media containing hormone. For imaging, discs were gently immobilized under a porous filter (Whatman cyclopore polycarbonate membranes, Sigma WHA70602513) in a 35mm glass-bottomed Petri dish (Mattek, Ashland, MA, USA; P35G-1.0-14-C) using double stick tape (Tesa 5338, doppelband fotostrip, ~100μm thick) as a spacer between the glass and the filter. To construct this chamber, we first punched a hole (6mm) in the tape, and then adhered the tape to the glass. The discs were transferred in ~10μl media to the center of the hole in the tape spacer. Care was taken to keep the tape dry. Discs were carefully arranged to be apical-side down (toward coverslip) using forceps. Most of the media was then removed, and the filter (cut approximately to the size of the tape) was quickly placed over the sample and firmly adhered to the tape with forceps. The dish was then filled with 2-3ml media. The height of these chambers is somewhat greater than the height of the discs, but the presence of the filter isolates the discs from flows and somewhat constrains disc movements so that they usually remain in the field of view. As an alternative, we also tried to immobilize discs using methylcellulose (Aldaz et al. 2010) dissolved in our culture medium, but discs do not proliferate in this condition.

During imaging, we used a syringe pump (PHD ULTRA, Harvard Apparatus, Cambridge, MA, USA) to automatically and continuously exchange the media during the course of imaging at a rate of 0.03 ml/min. The start of the movie was considered to be time 0hr. This time corresponds to 45-60min from the start of dissection (time required for sample preparation and microscope setup).

Imaging was performed using a Zeiss spinning disc microscope consisting of an AxioObserver inverted stand, motorized xyz stage with temperature control set to 25°C, Yokogawa CSU-X1 scanhead, and a Zeiss AxioCam MRm camera (2x2 binning), all controlled by Axiovision software. Discs were imaged through a Zeiss C-Apochromat 63X 1.2NA water immersion objective heated to 25°C with an objective heater. To capture the whole pouch, we acquired a 2x2 tiled region (10% overlap). Each region consisted of a Z-stack of 65-85 frames, spaced 0.5μm apart. Tiled Z-stacks were captured every 5min. We kept light exposure as low as possible to achieve a segmentable image. We used a power meter (PT9610, Gigahertz-optik, Munich, Germany) to measure the power of the laser through a 10x/0.45 NA objective within a week of the experiment. For imaging, we used a laser power of 0.04-0.05 mWatts and an exposure time of 350ms per image.

### Analysis of live imaging

#### Image processing, segmentation, and cell tracking

Raw data was first processed with a low-pass frequency filter to remove high frequency noise, followed by a rolling ball background subtraction algorithm using FIJI (Schindelin et al. 2012). The apical plane was projected onto 2D using a custom-made algorithm. We identified the two manifolds formed by the disc proper layer and the peripodial layer, respectively, in the 3D z-stacks, based on maximal brightness of the E-Cadherin-GFP signal and subject to hard constraints on the distance between the two layers and the slope of each manifold. We employed the algorithm of Wu and Chen (Wu & Chen 2002) to simultaneously determine the two manifolds that are optimal, i.e. as bright as possible under the given constraints. This algorithm and its application to the wing disc will be described in detail in a separate upcoming manuscript.

Projected tiles were stitched using the Grid/Collection Stitching FIJI plugin (Preibisch et al. 2009). Segmentation and cell tracking was performed on the projected images of the timelapse using Tissue Analyzer (Aigouy et al. 2016).

Cells within the wing pouch of the disc, as well as those on either side of the AP and DV boundaries, were manually marked using Tissue Miner (Etournay et al. 2016). The AP boundary is detectable on the basis of larger apical areas of the cells on either side, straighter cell boundaries at the interface and a slight reduction of E-Cadherin intensity on its anterior side. The DV boundary is detectable on the basis of increased E-Cadherin intensity and reduced apical areas of cells on either side of the boundary. All analysis was performed on the subset of cells within the pouch that were trackable throughout the entire course of the movie, excluding cells that move in or out of the field of view.

#### Analysis of cellular contributions to changes in tissue size and shape

The accumulated local tissue shear presented in Fig. 7A was obtained by dividing the tissue using a rectangular grid and calculating the tissue shear in each grid element as described previously (Etournay et al. 2016; Etournay et al. 2015). The blue bars represent the cumulative sum of tissue shear in each region, starting from 2hr (after the adaptation phase) and continuing to the end of the movie. For this calculation, we considered only grid elements with triangles covering more than half of the grid element area.

We quantify the relative area change rate and shear rate, averaged in the tracked wing pouch region, and the corresponding cellular contributions using the Triangle Method (Merkel et al. 2017). The relative area change decomposition states

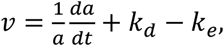

where *a* is the average cell area and *k_d_* and *k_e_* are cell division and extrusion rates. The relative area change rate *v* corresponds to the trace of the velocity gradient tensor *d_i_υ_j_*. The indices *i* and *j* take values *x* and *y*. In Fig. 7Bi, we plot the accumulated relative area change of the tracked region of the wing pouch 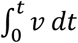 (blue line), accumulated relative change of the average cell area 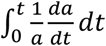 (green line), accumulated contribution due to cell divisions 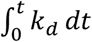 (orange line) and the accumulated contribution due to cell extrusions 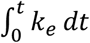 (cyan line).

The decomposition of the shear rate into its cellular contributions is

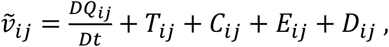

where *Q_ij_* is the cell elongation tensor and *D/Dt* is a corotational derivative (Merkel et al. 2017). *T_ij_*, *C_ij_*, *E_ij_* and *D_ij_* are contributions to the shear rate from T1 transitions, cell divisions, cell extrusions and correlation effects, respectively. In Fig. 7Bii we plot the accumulated xx component of shear of the tracked region 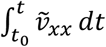 (blue line), accumulated xx component of shear due to cell elongation change 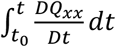 (green line), accumulated xx component of shear due to T1 transitions 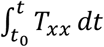 (red line), accumulated xx component of shear due to cell divisions 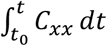 (orange line), accumulated xx component of shear due to cell extrusions 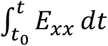 (cyan line) and accumulated xx component of shear due to correlation effects 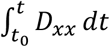 (magenta line). The x-axis of the coordinate system is parallel to the DV boundary as explained in the main text. The initial timepoint is *t*_0_ = 2hr.

The expansion of the wing pouch along the *x*-axis and the expansion along *y*-axis perpendicular to it are quantitatively described by velocity gradient components *∂_x_υ_x_* and *∂_y_υ_y_*, respectively. These components of the velocity gradient tensor are determined from *υ* and 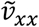 as

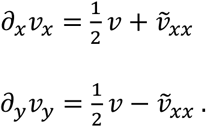

We then decompose the velocity gradient components into cellular contributions using the decompositions of *v* and 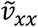 as follows

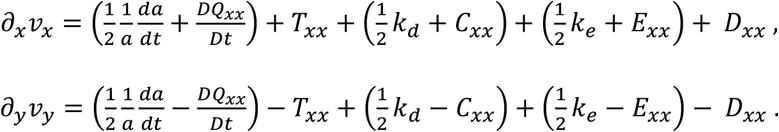

Here, 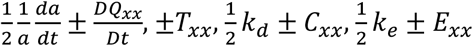 and ±*D_xx_* are the contributions to the velocity gradient component from cell shape change, T1 transitions, cell divisions, cell extrusions and correlations, respectively. The upper sign is used in the decomposition of *∂_x_υ_x_* and lower sign in the decomposition of *∂_y_υ_y_*.

In Fig. 7Biii we plot the accumulated wing pouch expansion along the x-axis 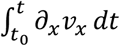 (blue line), accumulated cell shape expansion along the x-axis 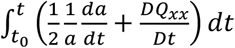 (green line), accumulated contribution to the expansion along the x-axis due to T1 transitions 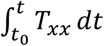 (red line), accumulated contribution to the expansion along the x-axis due to cell divisions 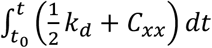 (orange line), accumulated contribution to the expansion along the x-axis due to cell extrusions 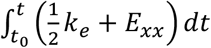 (cyan line) and accumulated contribution to the expansion along the x-axis due to correlation effects 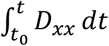 (magenta line).

In Fig. 7Biv, we plot the accumulated wing pouch expansion along the y-axis 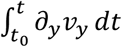 (blue line), accumulated cell shape expansion along the y-axis 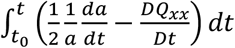 (green line), accumulated contribution to the expansion along the y-axis due to T1 transitions 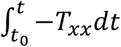 (red line), accumulated contribution to the expansion along the y-axis due to cell divisions 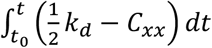 (orange line), accumulated contribution to the expansion along the y-axis due to cell extrusions 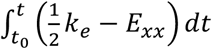 (cyan line) and accumulated contribution to the expansion along the y-axis due to correlation effects 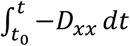 (magenta line).

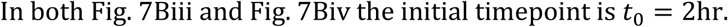

#### Estimation of tracking errors

Even if the cells are perfectly segmented in each frame of the timelapse, the automated cell tracking can still produce errors in tracking. These errors include false positive and false negative divisions, false positive extrusions and false positive cell appearances. The tracking errors either falsely remove or add a cell to the tissue. Since the overall cell number does not depend on the precision of tracking, the tracking errors always appear in pairs. For example, if a cell track is lost from frame 1 to 2, it is given a new cell ID in frame 2. This one error would cause the counting of a false extrusion (as the cell is thought to be lost in frame 1) and a false cell appearance (to account for the arrival of a new unique track in frame 2). Alternatively, the new cell ID in frame 2 could be attributed to a false cell division, if it is interpreted to be the daughter of a (false) division by a neighboring cell. These pairs of false cell events occur close to each other in space and time. We use this property to identify the tracking errors and adjust our quantification of the cellular contributions to tissue area change accordingly.

We have never observed any insertion of new cells into the wing disc epithelium and we thus assume that any cell appearance, not associated with a cell division, is due to an error in either segmentation or tracking. We devised a method to independently identify candidates for false positive and false negative divisions using the fact that just before the division, the apical area of the cell increases significantly as the nucleus moves apically. We measure the maximal cell area and the ratio of maximal cell area to the average of the cell area throughout the timelapse for each cell. We then discriminate dividing and non-dividing cells by performing a k-means classification on these two variables (Jones et al. 2001). One type of error that could not be identified by this method might occur if the original tracking identifies two cell divisions for the same cell where only one actually occurs. In these situations, when the divisions are separated by less than six frames, we take the later division to be true and the earlier one to be false. Using these candidates for dividing and non-dividing cells we construct a list of candidates for false positive and false negative divisions in the original tracking.

Finally, we pair cell appearances with cell extrusions and false negative cell division candidates, and then we pair the false positive cell division candidates with the remaining cell extrusions and remaining false negative cell division candidates. The cell events we manage to pair in this way we treat as false, and we adjust the cell event counting accordingly.

To construct pairs of these cell events where each event can appear at most once, we introduce a distance measure between cell events A and B in time and space:

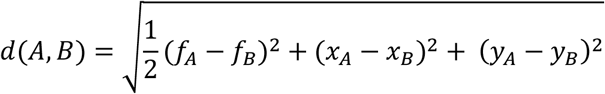

where *x_A,B_* and *y_A,B_* are coordinates of the cell centers (in pixels) in frame *f_A,B_* of the cell event, for each of the two cell events. We then iteratively construct pairs of false cell events from all possible combinations by repeating the following steps:

find a cell event pair with minimum d(A,B)

- if d(A, B) > 25 stop the procedure
- if d(A, B) < 25 identify pair (A, B) as a pair of false cell events
- remove the events A and B from consideration in future iterations

This method identifies more than 90% of the cell appearances as false. It also finds that between 85-90% of cell divisions and 45-75% of cell extrusions initially identified by Tissue Analyzer were true events and that between 10-20% of true divisions were not recognized by Tissue Analyzer. We used the corrected numbers of cell divisions and extrusions to calculate contributions to tissue area change. The remaining number of cell appearances is very low (<10) in all three experiments and not shown in the relative area change plots. This method does not identify the daughter cells. Therefore, in the shear calculation by the Triangle Method, we used the original tracking by the Tissue Analyzer. If we assume that the errors in cell division tracking are not correlated with the cell division orientation and that the falsely identified cell divisions are not oriented on average, we can estimate the relative error of the shear due to cell divisions to be equal to the fraction of true division events not recognized by the Tissue Analyzer i.e. 10-20%.

## ACKNOWLEDGEMENTS

RNA sequencing was performed by Annekathrin Kränkel and Andreas Dahl in the Deep Sequencing Group SFB655 at the Biotechnology Center of the Technische Universitaet Dresden. The processing of the RNAseq data was performed with considerable help from Holger Brandl of the Bioinformatics facility of the MPI-CBG. We thank Marko Brankatschk, Christian Dahmann, and Savraj Grewal for critical review of the manuscript prior to submission.

## Figure 1--Figure Supplement 1

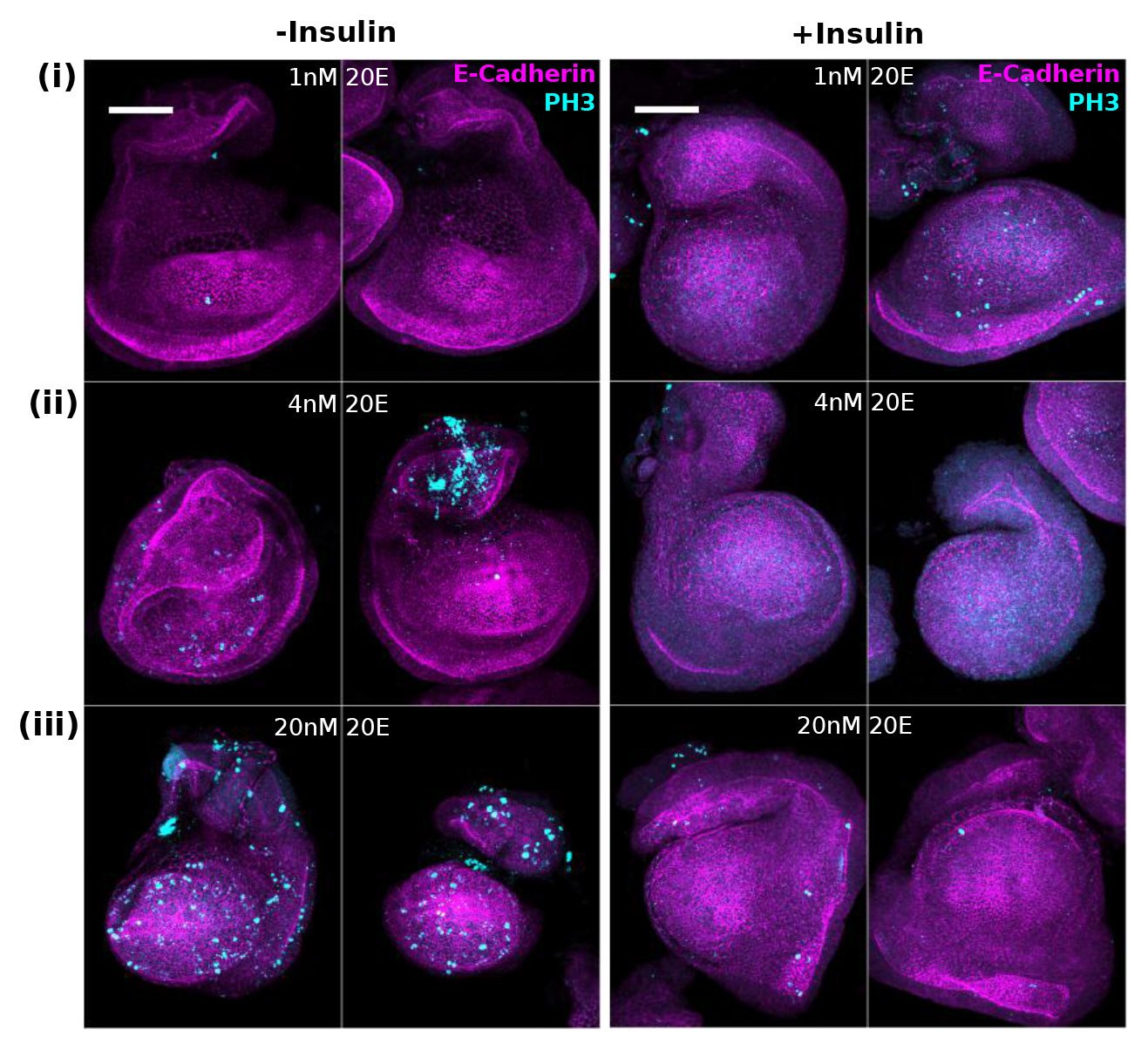
Effect of insulin on proliferation is not improved by lower 20E concentration. To determine if lower concentrations of 20E would allow insulin to further improve proliferation, discs were cultured for 24hr in 1nM (**i**), 4nM (**ii**) or 20nM (**iii**) 20E, with or without 5ug/ml insulin (right/left, respectively). After this length of culture, discs start to lose their morphology, making quantification difficult. In particular in insulin (+/−20E), the pouch region balloons forward and out. In all discs, the prospective notum starts to curl forward. Only in 20nM 20E alone (without insulin) is there any consistent division remaining. Best attempts were made to orient images with anterior to the left and dorsal up. Scale bar is 50um.

**Figure 3--Figure Supplement 1:**
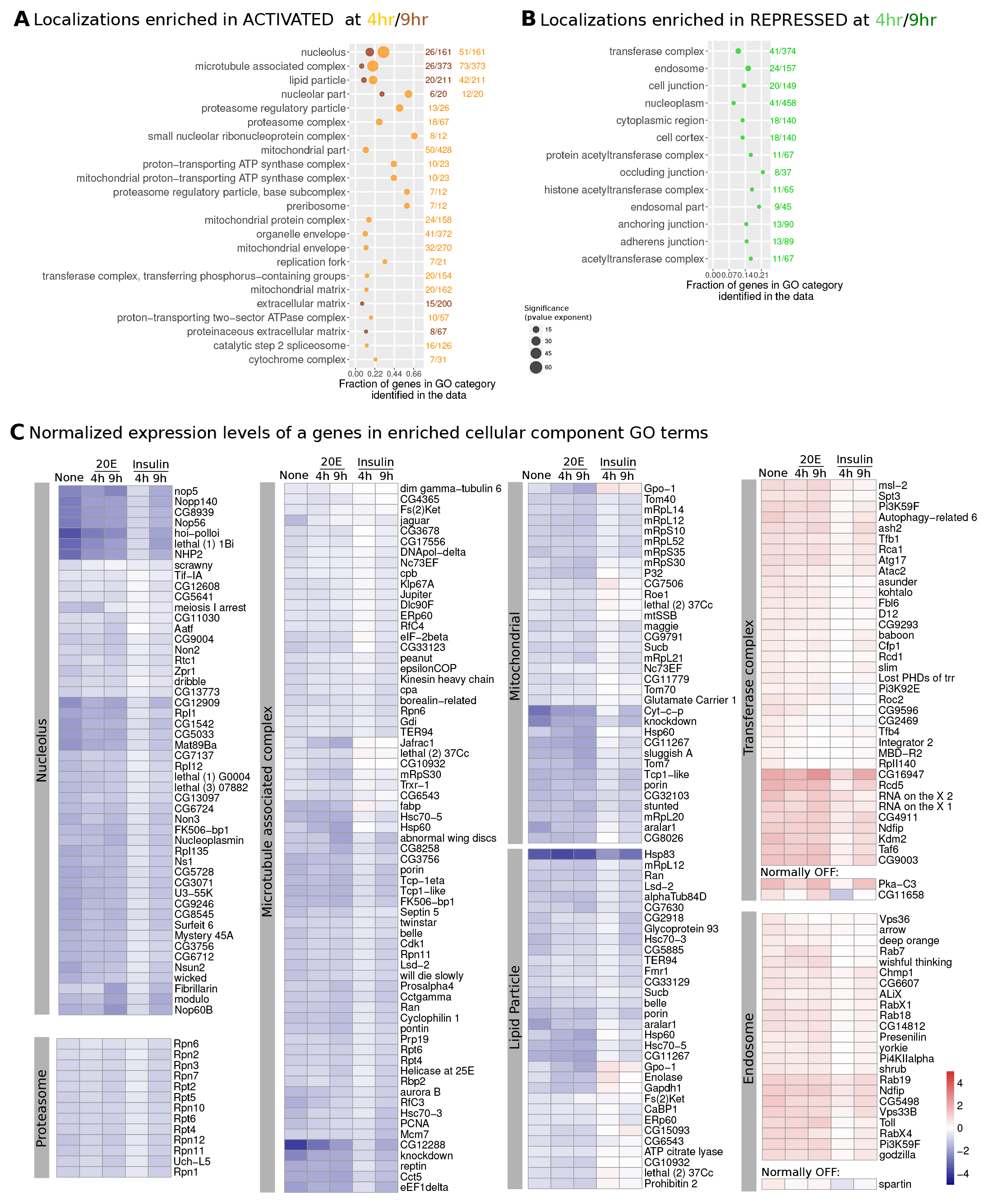
Cellular component GO terms enriched in insulin-regulated geneset. The insulin-regulated genes were analyzed as in Fig. 3 but using the Cellular Component (CC) terms. In both **A** and **B**, the fraction of total genes in the GO category represented in the dataset is plotted along x, with the actual numbers written on the right. The size of the point indicates the significance of enrichment (BH-corrected p-value). The cutoff for enrichment was a p-value <0.005. The color indicates the number of genes in that group at either 4hr of culture (light orange in **A**, green in **B**) or 9hr of culture (dark orange in **A**, dark green in **B**). (C) The expression of selected genes (fpkm) in related CC-enriched GO categories is shown for all of the tested conditions, normalized to its levels in uncultured wing discs (log 2(x/uncultured). In rare cases when a gene was ectopically activated (expression in culture but not in uncultured discs, labeled as “normally OFF”), its levels were normalized to the threshold for expression (5fpkm).

**Figure 3--Figure Supplement 2:**
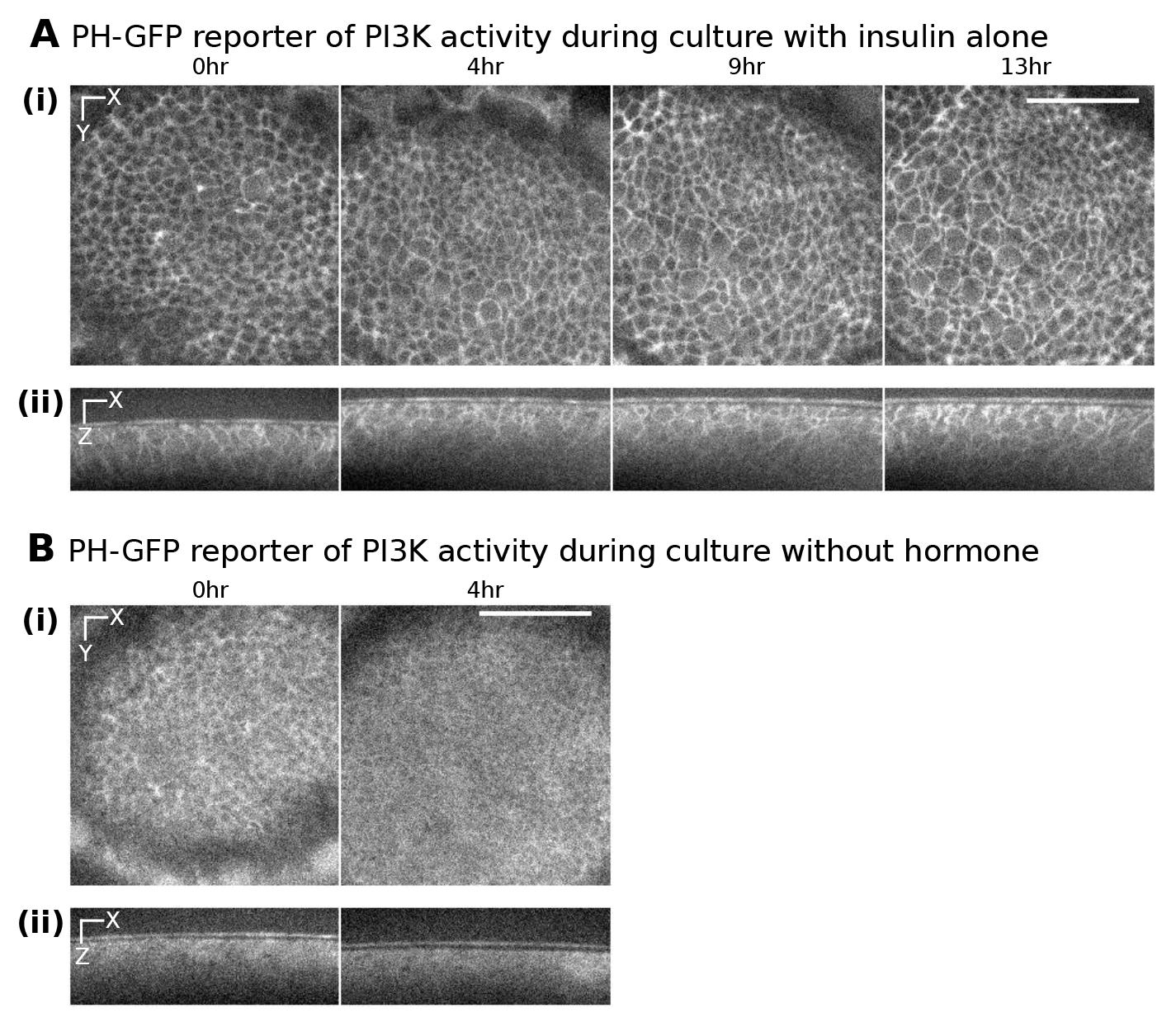
PI3K activity reporter remains active during long term culture in insulin. PH-GFP reporter of PI3K activity during culture with insulin (**A**) or no hormone (**B**). This reporter localizes to the membrane upon PIP3 production, thus reflecting PI3K activity. A single xy plane is shown in (**i**) and an xz slice through the middle of the disc is shown in (ii). Timepoints shown in each part are from the same disc grown on the microscope. Scale bar is 20um.

**Figure 4--Figure Supplement 1:**
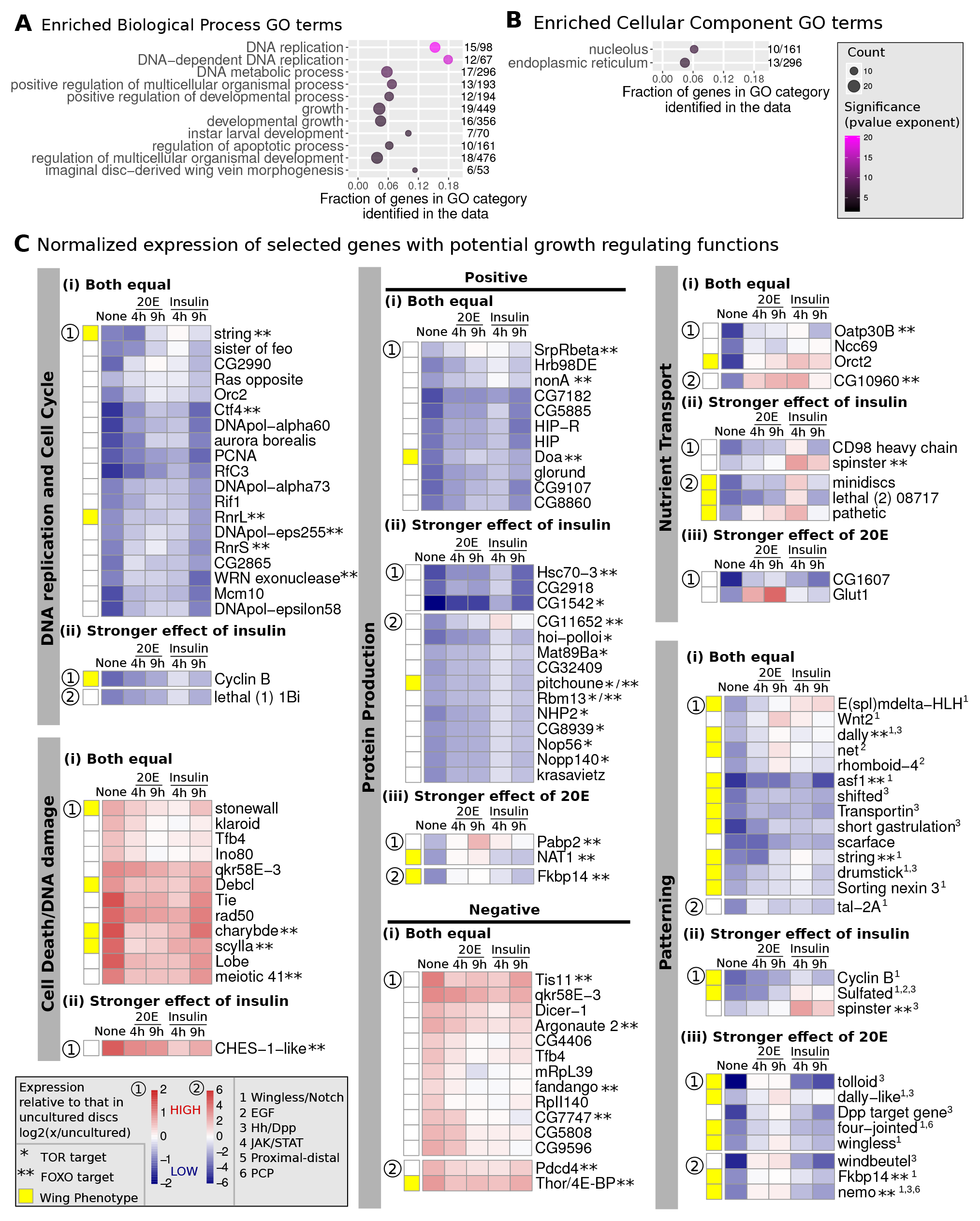
Partial overlap between insulin- and 20E-regulated genesets. Enriched biological process (**A**) or cellular component (**B**) GO terms were identified in the set of genes activated by both insulin (at 4hr) and 20E (at 9hr). A semantic similarity algorithm was used to reduce redundancy, with a similarity cutoff of 0.95. Labels on the right indicate the number of genes in the data over total genes in that GO category. (**C**) The expression of genes in selected functional categories (fpkm) is shown for all of the tested conditions, normalized to its levels in uncultured wing discs, log2(x/uncultured). Genes in each group were further subdivided based on whether the two hormones had equal (**i**) or unequal effects (**ii**, insulin stronger vs **iii**, 20E stronger). Genes were plotted on one of two scales (labeled as 1 or 2), depending on how much their values changed across conditions. Yellow squares indicate known phenotypes in the wing (at any stage of development) for loss of function for genes positively regulated by hormone or gain of function for genes negatively regulated by hormone. Asterisks indicate previously identified TOR (*) or FOXO (**) targets. Genes with known patterning functions (in any context) are indicated with numbers.

**Figure 6--Figure Supplement 1:**
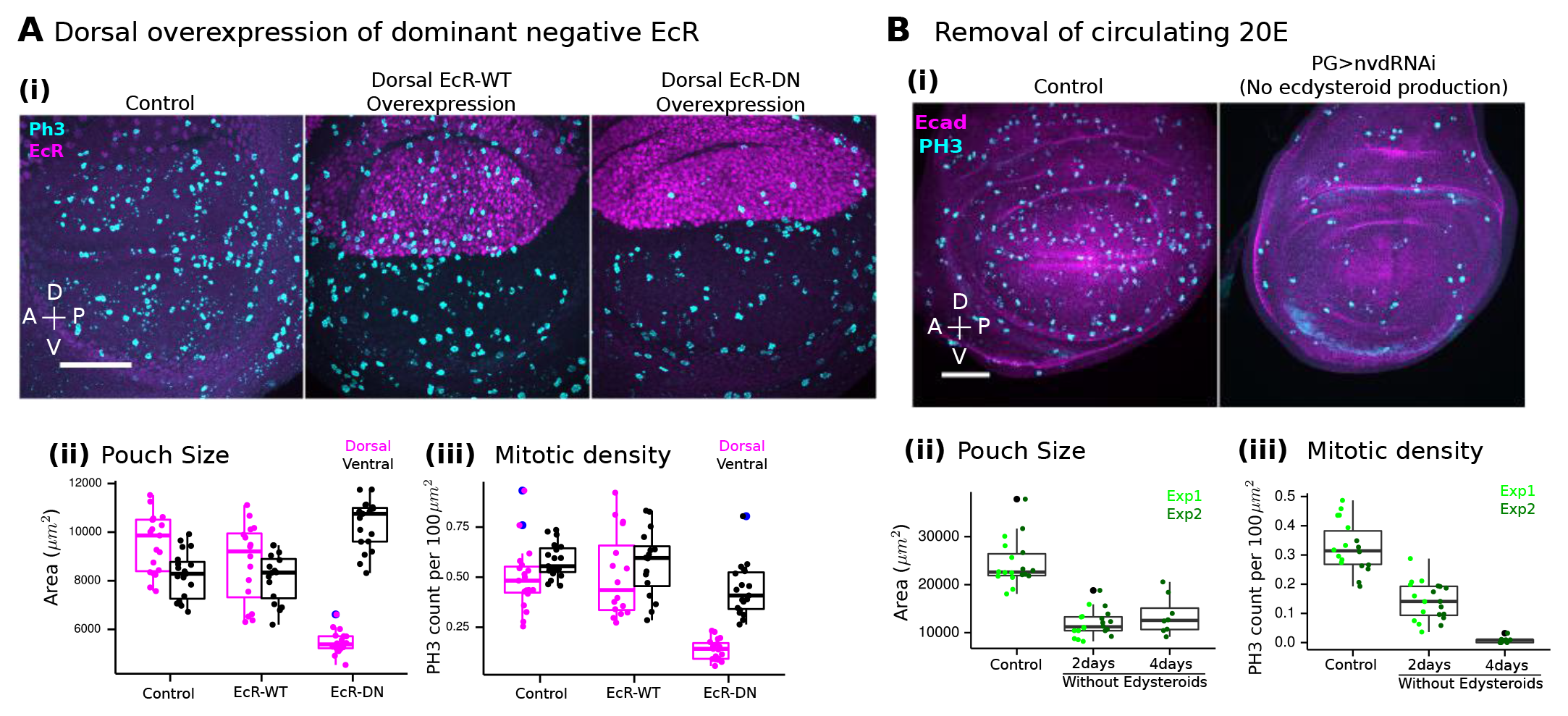
20E-signaling is autonomously required for wing growth during the 3rd instar. The effect of 20E on larval wing growth was verified by autonomously perturbing 20E signaling by overexpressing a dominant-negative EcR in the dorsal compartment of the wing (**A**) or by genetically preventing the synthesis of 20E (**B**). **A**. *Apterous-GAL4*, which is expressed in the dorsal compartment of the wing disc, combined with *tub>GAL80ts* was used to conditionally overexpress either a wild type (EcR-WT) or dominant negative EcR allele (EcRB1-W650A, called EcR-DN). Overexpression of the EcR construct was prevented in early developmental stages by growing the larvae at 18C, and then switching to 29C for 24hr during the third instar. The negative control is the *apterous-GAL4, gal80ts* crossed to wild type (left). Additionally, we verified that overexpression of wild type EcR (B1 isoform) would not affect growth (middle). (**ii**) Pouch size was quantified by measuring area in a maximum projection image. The Dorsal (pink) and Ventral (black) comparments were separately measured, estimating the boundary using E-Cadherin labeling. (**iii**) Density of PH3 staining in the Dorsal (pink) and Ventral (black) compartments. **B** *Phantom-GAL4*, which is expressed in the prothoracic ecdysteroid-producing gland of the brain (PG), combined with *tub>GAL80ts* was used to induce an RNAi targeting *neverland (nvd)*, a gene required to synthesize ecdysone. Induction of the RNAi was prevented in early developmental stages by growing the larvae at 18C, and then switching to 29C at the third instar. Control larvae contained only the *phantom-GAL4, tub>GAL80ts* crossed to wild type, and formed pupae after 2 days at 29C, whereas nvd-RNAi larvae stayed as larvae for several days. Disc size (area in a maximum projection image, **ii**) and density of mitotic nuclei (marked by phospho-histone H3, PH3, **iii**) were measured in the pouch region, defined as the region surrounded by the innermost folds. Data from two independent replicates of the experiment are separately colored. Only pupae remained for the control after 2 days at 29C; thus, only the 4-day timepoint is shown. For both **A** and **B**: (**i**) representative images show PH3 staining (cyan) as a maximum projection overlaid with either E-Cadherin (**A**) or EcR (**B**) in magenta. Dorsal is up and Anterior to the left. Scale bar is 50um. In (**ii**) and (**iii**), each dot represents one disc, with bars showing the extent of the first and third quartiles; lines reach up to 1.5* IQR (interquartile range); outliers outside of this range are blue (left) or black (right) dots.

**Figure 6--Figure Supplement 2:**
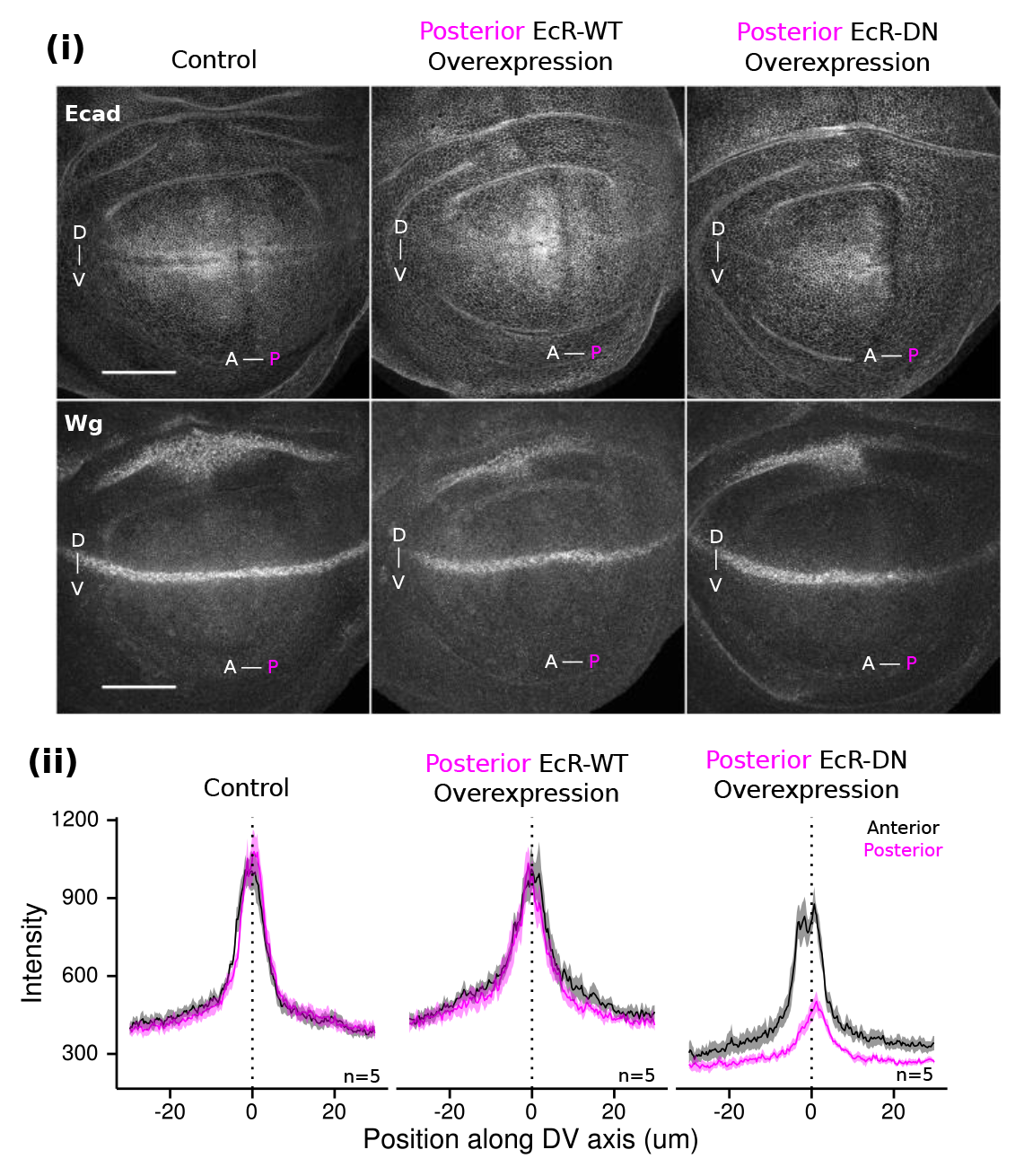
20E-signaling is required in the posterior compartment for Wg expression. The autonomous effect of 20E-signaling on Wg expression was verified using by overexpressing EcR-DN (EcRBl-W650A) in the posterior compartment. *Engrailed-GAL4*, which is active in the posterior wing disc, was combined with *tub>GAL80ts* to limit expression of the transgenes to 24hr in the third instar using a temperature shift. The control (left) was *engrailed-GAL4, GAL80ts* crossed to wild type flies. Additionally, it was verified that overexpression of a wild type EcR (EcRB1 isoform) did not affect Wg expression (middle) (**i**) Representative images of E-Cadherin (top) or Wg (bottom) staining in the same discs. Dorsal is up, anterior to the left. Markers deliniate the boundaries. Scale bars are 50um. (**ii**) Wg expression was quantified by measuring the absolute intensity alone a line drawn from Dorsal to Ventral of the pouch in the Anterior (black) or Posterior (pink) compartments. Posterior overexpression of EcR-DN (right) but not EcR-WT (middle) affects the Wg gradient. Control and perturbation stainings were done in parallel and imaged with the same acquisition settings on the same day.

**Figure 6--Figure Supplement 3:**
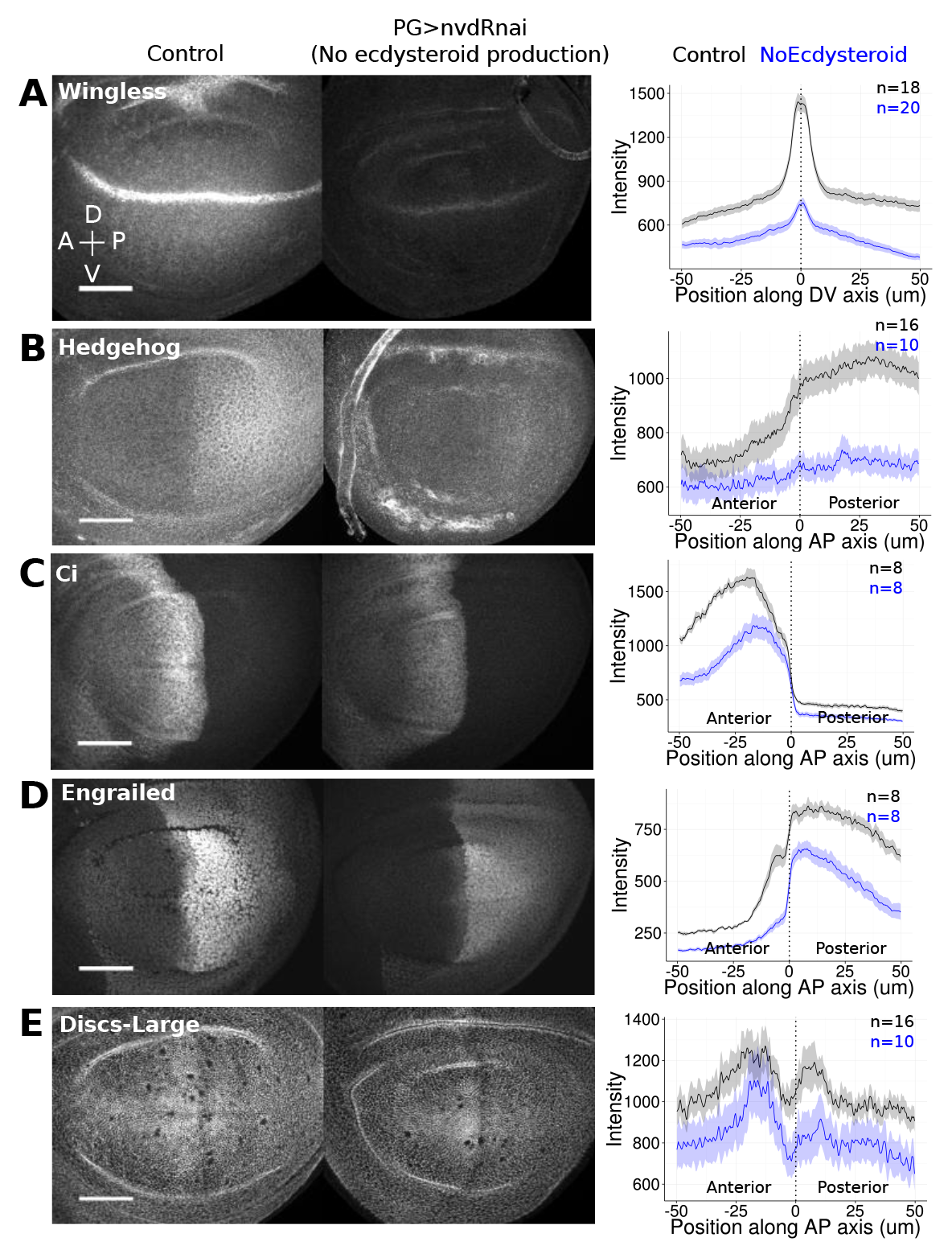
Circulating ecdysone is required for wing pattern during growth. *Phantom-GAL4*, which is expressed in the prothoracic ecdysteroid-producing gland of the brain (PG), combined with *tub>GAL80TS* was used to induce RNAi against *neverland* (nvd), a gene required to synthesize ecdysone, only during the third larval instar. Expression of Wingless (**A**), Hedgehog (**B**), Ci, the transcriptional activator downstream of Hh signaling (**C**), Engrailed, a Hh target gene (**D**), and Discs-large, a septate junction marker (**E**) was analyzed by immunofluorescence. Quantification of the changes in expression was performed by plotting the absolute intensity along a line drawn from one compartment to another. For (**A**), the line was drawn from Dorsal to Ventral in the Anterior comparment. For (**B**)-(**E**), the line was drawn from Anterior to Posterior in the Dorsal compartment. Black is the control; blue is the nvd-RNAi animals. The dark line is the mean for all discs measured in the same staining experiment; the shaded ribbon indicates the standard deviation. Control and perturbation stainings were done in parallel and imaged with the same acquisition settings on the same day. Scale bars correspond to 50um.

## REFERENCES

Aegerter-Wilmsen, T. et al., 2012. Integrating force-sensing and signaling pathways in a model for the regulation of wing imaginal disc size. Development (Cambridge, England), 139(17), pp.3221–31.

Affolter, M. & Basler, K., 2007. The Decapentaplegic morphogen gradient: from pattern formation to growth regulation. Nature reviews. Genetics, 8(9), pp.663–674.

Aigouy, B., Umetsu, D. & Eaton, S., 2016. Segmentation and Quantitative Analysis of Epithelial Tissues. In pp. 227–239.

Aldaz, S., Escudero, L.M. & Freeman, M., 2010. Live imaging of Drosophila imaginal disc development. Proceedings of the National Academy of Sciences of the United States of America, 107(32), pp.14217–22.

Alic, N. et al., 2014. Interplay of dFOXO and Two ETS-Family Transcription Factors Determines Lifespan in Drosophila melanogaster S. K. Kim, ed. PLoS Genetics, 10(9), p.e1004619.

Ambegaonkar, A.A. et al., 2012. Propagation of dachsous-fat planar cell polarity. Current Biology, 22(14), pp.1302–1308.

Andres, A.J. & Thummel, C.S., 1994. Methods for quantitative analysis of transcription in larvae and prepupae. Methods in cell biology, 44, pp.565–73.

Arganda-Carreras, I. et al., 2017. Trainable Weka Segmentation: a machine learning tool for microscopy pixel classification. Bioinformatics, 9, pp.676–682.

Arquier, N. et al., 2008. Drosophila ALS Regulates Growth and Metabolism through Functional Interaction with Insulin-Like Peptides. Cell Metabolism, 7(4), pp.333–338.

Aw, W.Y. & Devenport, D., 2016. Planar cell polarity: Global inputs establishing cellular asymmetry. Current Opinion in Cell Biology.

Ayers, K.L. et al., 2012. Dally and Notum regulate the switch between low and high level Hedgehog pathway signalling. Development, 139(17), pp.3168–3179.

Ayers, K.L. et al., 2010. The Long-Range Activity of Hedgehog Is Regulated in the Apical Extracellular Space by the Glypican Dally and the Hydrolase Notum. Developmental Cell, 18(4), pp.605–620.

Baena-López, L.A., Baonza, A. & García-Bellido, A., 2005. The orientation of cell divisions determines the shape of Drosophila organs. Current biology: CB, 15(18), pp.1640–4.

Balaji, R. et al., 2017. Calcium spikes, waves and oscillations in a large, patterned epithelial tissue. Scientific Reports, 7, p.42786.

Beckstead, R.B., Lam, G. & Thummel, C.S., 2005. The genomic response to 20-hydroxyecdysone at the onset of Drosophila metamorphosis. Genome Biology, 6(12), p.R99.

Beira, J. V. & Paro, R., 2016. The legacy of Drosophila imaginal discs. Chromosoma, 125(4), pp.573–592.

Bodenstein, D., 1943. Hormones and tissue competence in the development of Drosophila. The Biological Bulletin, 84(1), pp.34–58.

Bornemann, D.J. et al., 2004. Abrogation of heparan sulfate synthesis in Drosophila disrupts the Wingless, Hedgehog and Decapentaplegic signaling pathways. Development (Cambridge, England), 131(9), pp.1927–38.

Brennan, C.A. et al., 2001. Broad-complex, but not ecdysone receptor, is required for progression of the morphogenetic furrow in the Drosophila eye. Development (Cambridge, England), 128(1), pp.1–11.

Brennan, C.A. et al., 1998. Ecdysone pathway is required for furrow progression in the developing Drosophila eye. Development (Cambridge, England), 125(14), pp.2653–64.

Brittle, A., Thomas, C. & Strutt, D., 2012. Planar polarity specification through asymmetric subcellular localization of fat and dachsous. Current Biology, 22(10), pp.907–914.

Britton, J.S. & Edgar, B.A., 1998. Environmental control of the cell cycle in Drosophila: nutrition activates mitotic and endoreplicative cells by distinct mechanisms. Development, 125(11).

Bryant, P.J. et al., 1988. Mutations at the fat locus interfere with cell proliferation control and epithelial morphogenesis in Drosophila. Developmental Biology, 129(2), pp.541–554.

Bryant, P.J. & Levinson, P., 1985. Intrinsic growth control in the imaginal primordia of Drosophila, and the autonomous action of a lethal mutation causing overgrowth. Developmental Biology, 107(2), pp.355–363.

de Celis, J.F. & Bray, S., 1997. Feed-back mechanisms affecting Notch activation at the dorsoventral boundary in the Drosophila wing. Development (Cambridge, England), 124(17), pp.3241–3251.

Chen, H. & Boutros, P.C., 2011. VennDiagram: a package for the generation of highly-customizable Venn and Euler diagrams in R. BMC bioinformatics, 12(1), p.35.

Cheng, L.Y. et al., 2011. Anaplastic lymphoma kinase spares organ growth during nutrient restriction in drosophila. Cell, 146(3), pp.435–447.

Clark, H.F. et al., 1995. Dachsous encodes a member of the cadherin superfamily that controls imaginai disc morphogenesis in Drosophila. Genes and Development, 9(12), pp.1530–1542.

Delanoue, R., Slaidina, M. & Léopold, P., 2010. The Steroid Hormone Ecdysone Controls Systemic Growth by Repressing dMyc Function in Drosophila Fat Cells. Developmental Cell, 18(6), pp.1012–1021.

DeVido, S.K. et al., 2008. The role of Polycomb-group response elements in regulation of engrailed transcription in Drosophila. Development (Cambridge, England), 135(4), pp.669–676.

Etournay, R. et al., 2015. Interplay of cell dynamics and epithelial tension during morphogenesis of the Drosophila pupal wing. eLife, 4, p.e07090.

Etournay, R. et al., 2016. TissueMiner: A multiscale analysis toolkit to quantify how cellular processes create tissue dynamics. eLife, 5.

Eugster, C. et al., 2007. Lipoprotein-Heparan Sulfate Interactions in the Hh Pathway. Developmental Cell, 13(1), pp.57–71.

Fristrom, J.W., Logan, W.R. & Murphy, C., 1973. The synthetic and minimal culture requirements for evagination of imaginal discs of Drosophila melanogaster in vitro. Developmental Biology, 33(2), pp.441–456.

Garcia-Bellido, A. & Merriam, J.R., 1971. Parameters of the wing imaginal discdevelopment ofDrosophila melanogaster. Developmental Biology, 24(1), pp.61–87.

Gauhar, Z. et al., 2009. Genomic mapping of binding regions for the Ecdysone receptor protein complex. Genome research, 19(6), pp.1006–13.

Géminard, C., Rulifson, E.J. & Léopold, P., 2009. Remote Control of Insulin Secretion by Fat Cells in Drosophila. Cell Metabolism, 10(3), pp.199–207.

Gershman, B. et al., 2007. High-resolution dynamics of the transcriptional response to nutrition in Drosophila: a key role for dFOXO. Physiological genomics, 29(1), pp.24–34.

Gibson, M.C. et al., 2006. The emergence of geometric order in proliferating metazoan epithelia. Nature, 442(7106), pp.1038–1041.

González-Gaitán, M., Capdevila, M.P. & García-Bellido, A., 1994. Cell proliferation patterns in the wing imaginal disc of Drosophila. Mechanisms of Development, 46(3), pp.183–200.

Guertin, D.A. et al., 2006. Functional Genomics Identifies TOR-Regulated Genes that Control Growth and Division. Current Biology, 16(10), pp.958–970.

Guirao, B. et al., 2015. Unified quantitative characterization of epithelial tissue development. eLife, 4(DECEMBER 2015).

Hacker, U., Lin, X. & Perrimon, N., 1997. The Drosophila sugarless gene modulates Wingless signaling and encodes an enzyme involved in polysaccharide biosynthesis. Development, 124(18), pp.3565–3573.

Handke, B. et al., 2014. Towards long term cultivation of Drosophila wing imaginal discs in vitro. PloS one, 9(9), p.e107333.

Hartl, T.A. & Scott, M.P., 2014. Wing tips: The wing disc as a platform for studying Hedgehog signaling. Methods, 68(1), pp.199–206.

Heemskerk, I., Lecuit, T. & LeGoff, L., 2014. Dynamic clonal analysis based on chronic in vivo imaging allows multiscale quantification of growth in the Drosophila wing disc. Development, 141(11).

Heller, D. et al., 2016. EpiTools: An Open-Source Image Analysis Toolkit for Quantifying Epithelial Growth Dynamics. Developmental Cell, 36(1), pp.103–116.

Herboso, L. et al., 2015. Ecdysone promotes growth of imaginal discs through the regulation of Thor in D. melanogaster. Scientific Reports, 5(1), p.12383.

Honegger, B. et al., 2008. Imp-L2, a putative homolog of vertebrate IGF-binding protein 7, counteracts insulin signaling in Drosophila and is essential for starvation resistance. Journal of biology, 7(3), p.10.

Huang, J. et al., 2009. Directed, efficient, and versatile modifications of the Drosophila genome by genomic engineering. Proceedings of the National Academy of Sciences, 106(20), pp.8284–8289.

Jaiswal, M., Agrawal, N. & Sinha, P., 2006. Fat and Wingless signaling oppositely regulate epithelial cell-cell adhesion and distal wing development in Drosophila. Development (Cambridge, England), 133(5), pp.925–935.

Jones, E., Oliphant, T. & Peterson, P., 2001. SciPy: Open Source Scientific Tools for Python. Available at: http://www.scipy.org/.

Kim, D. et al., 2013. TopHat2: accurate alignment of transcriptomes in the presence of insertions, deletions and gene fusions. Genome Biology, 14(4), p.R36.

Kozlova, T. & Thummel, C.S., 2000. Steroid regulation of postembryonic development and reproduction in drosophila. Trends in Endocrinology and Metabolism, 11(7), pp.276–280.

Lavrynenko, O. et al., 2015. The ecdysteroidome of Drosophila: influence of diet and development. Development, 142(21).

Legoff, L. et al., 2013. A global pattern of mechanical stress polarizes cell divisions and cell shape in the growing Drosophila wing disc. Development (Cambridge, England), 140(19), pp.4051–9.

Li, L. et al., 2010. Nutritional control of gene expression in Drosophila larvae via TOR, Myc and a novel cis-regulatory element. BMC Cell Biology, 11(1), p.7.

Mao, Y. et al., 2006. Dachs: an unconventional myosin that functions downstream of Fat to regulate growth, affinity and gene expression in Drosophila. Development (Cambridge, England), 133(13), pp.2539–51.

Mao, Y. et al., 2013. Differential proliferation rates generate patterns of mechanical tension that orient tissue growth. The EMBO Journal, 32(21), pp.2790–2803.

Mao, Y. et al., 2011. Planar polarization of the atypical myosin Dachs orients cell divisions in Drosophila. Genes & Development, 25(2), pp.131–136.

Marois, E., Mahmoud, A. & Eaton, S., 2006. The endocytic pathway and formation of the Wingless morphogen gradient. Development, 133(2).

Martin, M., 2011. Cutadapt removes adapter sequences from high-throughput sequencing reads. EMBnet.journal, 17(1), p.10.

Merkel, M. et al., 2014. The balance of prickle/spiny-legs isoforms controls the amount of coupling between core and fat PCP systems. Current Biology, 24(18), pp.2111–2123.

Merkel, M. et al., 2017. Triangles bridge the scales: Quantifying cellular contributions to tissue deformation. Physical Review E, 95(3), p.32401.

Micchelli, C.A. & Blair, S.S., 1999. Dorsoventral lineage restriction in wing imaginal discs requires Notch. Nature, 401(6752), pp.473–476.

Micchelli, C. a, Rulifson, E.J. & Blair, S.S., 1997. The function and regulation of cut expression on the wing margin of Drosophila: Notch, Wingless and a dominant negative role for Delta and Serrate. Development (Cambridge, England), 124, pp.1485–1495.

Milán, M., Campuzano, S. & García-Bellido, a, 1996. Cell cycling and patterned cell proliferation in the Drosophila wing during metamorphosis. Proceedings of the National Academy of Sciences of the United States of America, 93(21), pp.11687–92.

Milner, M.J., 1977. The eversion and differentiation of Drosophila melanogaster leg and wing imaginal discs cultured in vitro with an optimal concentration of β-ecdysone. Development, 37(1).

Mirth, C.K., Truman, J.W. & Riddiford, L.M., 2009. The ecdysone receptor controls the post-critical weight switch to nutrition-independent differentiation in Drosophila wing imaginal discs. Development (Cambridge, England), 136(14), pp.2345–53.

Mitchell, N.C. et al., 2013. The Ecdysone receptor constrains wingless expression to pattern cell cycle across the Drosophila wing margin in a cyclin B-dependent manner. BMC Developmental Biology, 13(1), p.28.

Okamoto, N. et al., 2013. A secreted decoy of InR antagonizes insulin/IGF signaling to restrict body growth in drosophila. Genes and Development, 27(1), pp.87–97.

Ono, H. et al., 2006. Spook and Spookier code for stage-specific components of the ecdysone biosynthetic pathway in Diptera. Developmental Biology, 298(2), pp.555–570.

Pastor-Pareja, J.C. & Xu, T., 2011. Shaping Cells and Organs in Drosophila by Opposing Roles of Fat Body-Secreted Collagen IV and Perlecan. Developmental cell, 21(2), pp.245–56.

Preibisch, S., Saalfeld, S. & Tomancak, P., 2009. Globally optimal stitching of tiled 3D microscopic image acquisitions. Bioinformatics (Oxford, England), 25(11), pp.1463–5.

Puig, O. et al., 2003. Control of cell number by Drosophila FOXO: downstream and feedback regulation of the insulin receptor pathway. Genes & development, 17(16), pp.2006–20.

Resino, J., Salama-Cohen, P. & García-Bellido, A., 2002. Determining the role of patterned cell proliferation in the shape and size of the Drosophila wing. Proceedings of the National Academy of Sciences of the United States of America, 99(11), pp.7502–7.

Rogulja, D., Rauskolb, C. & Irvine, K.D., 2008. Morphogen Control of Wing Growth through the Fat Signaling Pathway. Developmental Cell, 15(2), pp.309–321.

Rulifson, E.J. et al., 1996. wingless refines its own expression domian on the Drosophila wing margin. Nature, 384(6604), pp.72–4.

Rulifson, E.J., Kim, S.K. & Nusse, R., 2002. Ablation of Insulin-Producing Neurons in Flies: Growth and Diabetic Phenotypes. Science, 296(5570).

Sagner, A. et al., 2012. Establishment of global patterns of planar polarity during growth of the drosophila wing epithelium. Current Biology, 22(14), pp.1296–1301.

Schindelin, J. et al., 2012. Fiji: an open-source platform for biological-image analysis. Nature methods, 9(7), pp.676–82.

Schwank, G. et al., 2011. Antagonistic growth regulation by Dpp and Fat drives uniform cell proliferation. Developmental cell, 20(1), pp.123–30.

Spellberg, M.J., Marr, M.T. & II, 2015. FOXO regulates RNA interference in Drosophila and protects from RNA virus infection. Proceedings of the National Academy of Sciences of the United States of America, 112(47), pp.14587–92.

Teleman, A.A. et al., 2008. Nutritional Control of Protein Biosynthetic Capacity by Insulin via Myc in Drosophila. Cell Metabolism, 7(1), pp.21–32.

Trapnell, C. et al., 2012. Differential gene and transcript expression analysis of RNA-seq experiments with TopHat and Cufflinks. Nature Protocols, 7(3), pp.562–578.

Tsao, C.-K. et al., 2016. Long Term Ex Vivo Culture and Live Imaging of Drosophila Larval Imaginal Discs A. Bergmann, ed. PLOS ONE, 11(9), p.e0163744.

Waddington, C.H., 1940. The genetic control of wing development in Drosophila. Journal of Genetics, 41, pp.75–139.

Worley, M.I., Setiawan, L. & Hariharan, I.K., 2013. TIE-DYE: a combinatorial marking system to visualize and genetically manipulate clones during development in Drosophila melanogaster. Development, 140(15).

Wu, X. & Chen, D.Z., 2002. Optimal Net Surface Problems with Applications. In Springer, Berlin, Heidelberg, pp. 1029–1042.

Yan, D. & Lin, X., 2009. Shaping morphogen gradients by proteoglycans. Cold Spring Harbor perspectives in biology, 1(3).

Yoshiyama, T. et al., 2006. Neverland is an evolutionally conserved Rieske-domain protein that is essential for ecdysone synthesis and insect growth. Development, 133(13).

Yu, G. et al., 2012. clusterProfiler: an R Package for Comparing Biological Themes Among Gene Clusters. OMICS: A Journal of Integrative Biology, 16(5), pp.284–287.

Zartman, J., Restrepo, S. & Basler, K., 2013. A high-throughput template for optimizing Drosophila organ culture with response-surface methods. Development, 140(3).

